# A quantitative redox proteome of the human muscle response to exercise

**DOI:** 10.64898/2026.09.15.751463

**Authors:** Yu Lei, Jonathan J. Petrocelli, Anita Reddy, Carlos Henríquez-Olguin, Nils Burger, Sanghee Shin, Bingsen Zhang, Taylor Covington, Markus Waldeck-Weiermair, Jiuchun Zhang, Shelley M. Wei, Thomas E. Jensen, Christian T. Voldstedlund, Christian S. Carl, Bente Kiens, Erik A. Richter, Edward T. Chouchani

**Affiliations:** Department of Cancer Biology, Dana–Farber Cancer Institute, Boston, MA, USA; Department of Cell Biology, Harvard Medical School, Boston, MA, USA; Department of Nutrition, Exercise and Sports, The August Krogh Section for Human and Molecular Physiology, University of Copenhagen, Copenhagen, Denmark; Center for Exercise Physiology and Metabolism, Department of Kinesiology, Faculty of Medicine, Universidad Finis Terrae, Santiago, Chile; Heart and Vascular Institute, Mass General Brigham, Harvard Medical School, Boston, MA, USA; Gottfried Schatz Research Center, Molecular Biology and Biochemistry, Medical University of Graz, Graz, Austria; Howard Hughes Medical Institute, Chevy Chase, MD, USA

**Author notes:** These authors contributed equally (Yu Lei, Jonathan J. Petrocelli).

## Abstract

Reactive oxygen species (ROS) regulate protein function through reversible cysteine oxidation. In human skeletal muscle, exercise-induced ROS initiates adaptations such as mitochondrial biogenesis, increased insulin sensitivity, and hypertrophy. However, specific protein targets of ROS regulation during exercise remain unclear owing to longstanding challenges in analyzing redox proteomes *in vivo*. We applied cysteine derivatization and multiplexed proteomics to map muscle protein cysteine oxidation in humans during exercise. The OxiMuscle dataset quantifies reversible modifications across 9,177 unique cysteine sites on 2,782 proteins, comprising 17,492 individual cysteine site measurements in young men undergoing three types of exercise, providing the first comprehensive, site-resolved and quantitative analysis of the exercise-regulated redox cysteine proteome in humans. We systematically define cysteine oxidation targets regulated by at least one form of exercise, many of which reside in proteins with established roles in muscle physiology. Among these sites is a redox-regulated cysteine on the autophagy receptor protein p62. We demonstrate that reversible oxidation of this cysteine regulates p62-mediated autophagy upon myotube contraction and mouse muscle adaptation to exercise *in vivo*. Together, these results define a redox-driven mechanism linking exercise-induced autophagy to muscle adaptation. More broadly, our findings offer a comprehensive resource on redox-signaling networks in human muscle, accessible at http://oximuscle-alb-1899330623.us-east-1.elb.amazonaws.com/.

---

Redox reactions are fundamental biochemical processes that can facilitate communication in biological systems. This communication is mediated in part through the generation of reactive oxygen species (ROS), which can alter protein function through covalent modification of cysteine residues^1–3^. Redox modification of protein cysteines has emerged as a critical determinant of diverse biological outcomes. In the context of skeletal muscle physiology, ROS play a pivotal role in mediating adaptive responses to exercise^4–6^. Acute exercise induces a transient increase in muscle ROS levels, which is essential for initiating signaling that leads to beneficial adaptations. Antioxidant supplementation impairs exercise-mediated enhancements in mitochondrial biogenesis and insulin sensitivity in both humans and rodents^4,5,7,8^. Moreover, blunting the exercise-induced rise in muscle ROS with antioxidant supplementation can attenuate gains in lean mass during strength training, although the literature is mixed^9,10^. In particular, cytosolic ROS production by NADPH oxidase 2 has been shown to regulate muscle glucose uptake during exercise^11^. Furthermore, deleting NADPH oxidase 4 in mouse skeletal muscle accelerates the aging associated physiological decline, resulting in overt sarcopenia, frailty and insulin resistance^12,13^. Together, there is a general view that ROS are crucial for facilitating muscle improvements associated with exercise.

The established role of ROS signaling in skeletal muscle adaptation raises fundamental mechanistic questions. Foremost among these concerns the identity of the direct molecular targets through which ROS regulate muscle biology. To date, technical challenges associated with studying protein redox regulation *in vivo*^14^ have hindered the achievement of a comprehensive quantitative analysis of the redox proteome within muscle during exercise.

We recently developed a cysteine derivatization and enrichment method coupled with multiplexed mass spectrometry (MS) allowing for deep and quantitative analysis protein cysteine redox modifications in living tissues^15^. Here, we deployed this technology to comprehensively map muscle protein cysteine oxidation *in vivo* upon exercise in humans. This OxiMuscle dataset tracks the % reversible modification of ∼17,492 cysteine sites in human muscle of young men subjected to three forms of exercise, each known to remodel muscle tissue. This landscape represents the largest and, to our knowledge, first stoichiometrically resolved analysis of the exercise-regulated cysteine proteome in human muscle.

From this dataset, we observe cysteine oxidation targets that are shared across all forms of muscle exercise, as well as those specific to particular forms of exercise. We systematically categorize these targets to identify a range of biological processes subject to redox regulation upon exercise. Among the cysteines that undergo pronounced redox modification induced by exercise is Cys113 on the pivotal autophagy protein p62. We find that reversible oxidation of this cysteine plays a critical role in p62-mediated autophagosome formation and autophagic flux in muscle cells. Loss of p62 Cys113 significantly diminishes autophagosome formation induced upon chemogenomic production of H_2_O_2_ or myotube contraction, and in mice alters proteomic remodeling initiated in muscle following exercise. Together, we describe a redox mechanism that regulates exercise-induced autophagy and muscle adaptation to exercise *in vivo*. Moreover, these findings provide a comprehensive analysis of redox-signaling networks in human muscle, which can be accessed through an interactive web resource at http://oximuscle-alb-1899330623.us-east-1.elb.amazonaws.com/.

## RESULTS

### A Quantitative Landscape of Protein Cysteine Oxidation in Human Muscle Upon Exercise

Despite the importance of ROS signaling in exercising muscle, the specific modifications that underpin these mechanisms remain largely undefined^16^. Prior redox-proteomic surveys of muscle have focused on small numbers of pre-defined targets, or used non-stoichiometric methods of quantification^17,18^. Measuring site occupancy, defined as the extent of reversible modification at individual cysteine residues, is key to identifying functional nodes that enable ROS and related redox species to exert physiological functions. Although some proteomic techniques assess the stoichiometry of cysteine modifications, they typically suffer from low proteome coverage^19–22^. To overcome this limitation, we recently developed a method based on cysteine-reactive phosphate tags (CPT-MS), which combines cysteine derivatization and enrichment with multiplexed proteomic mass spectrometry^15,23^. CPT-MS enables comprehensive characterization of protein cysteine oxidation *in vivo*, simultaneously quantifying reversible oxidation states at tens of thousands of cysteine residues in a single experiment **(Figure S1A**; **Figure 1A)**.

**Figure 1.**
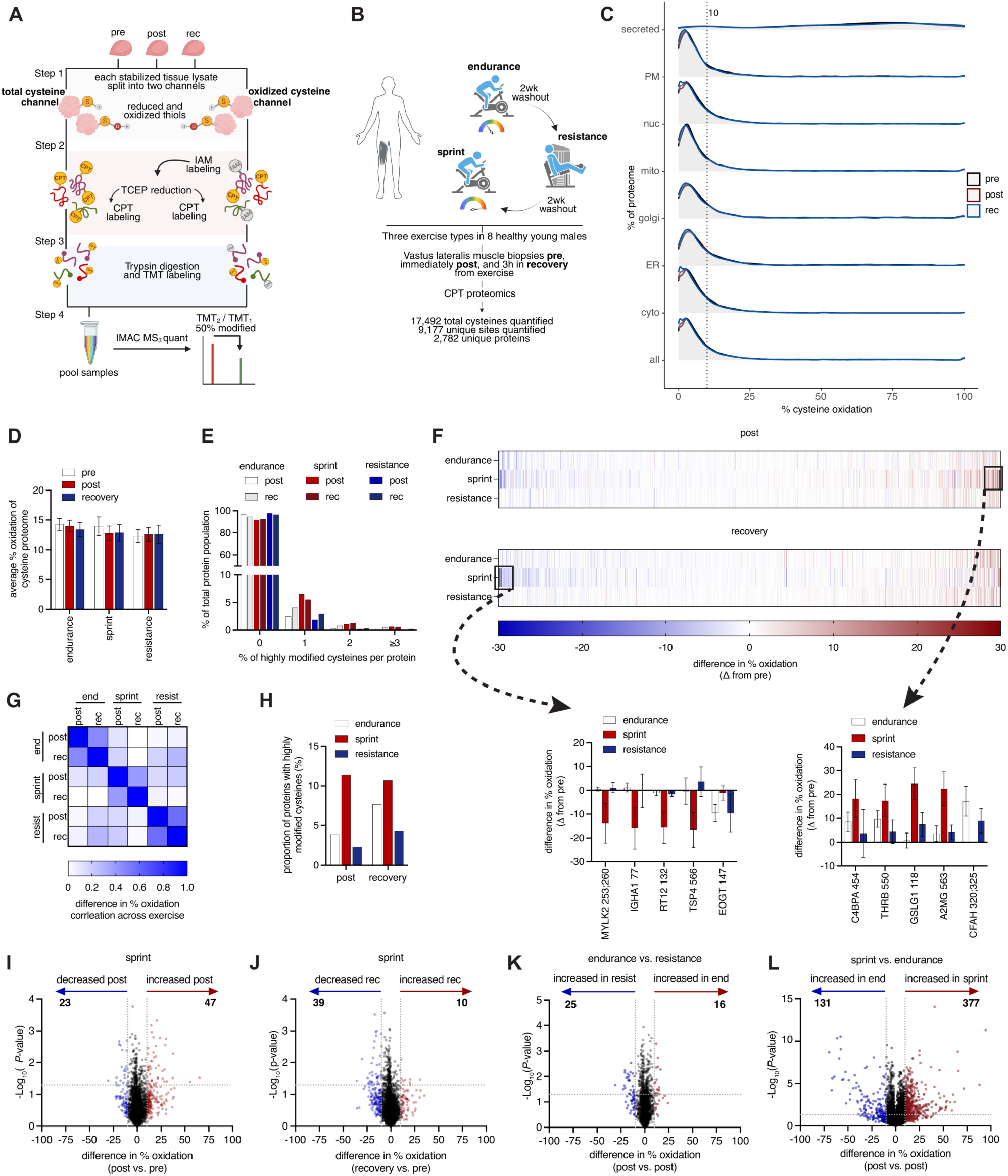
OxiMuscle Provides a Landscape of Redox Modification in Response to Acute Exercise in Human Skeletal Muscle. **(A)** Workflow for determining percent reversible modification of cysteines in human muscle pre, post, and after 3 hours of recovery from exercise using CPT. CPT, cysteine-reactive phosphate tag; IAM, iodoacetamide; rec, recovery. **(B)** Overview of OxiMuscle. Vastus lateralis muscle was collected from 8 healthy young men pre, post and in recovery from an acute bout of endurance exercise. The same males underwent a 2-week washout period of no exercise followed by an acute bout of resistance exercise, then another 2-week washout period and an acute bout of sprint exercise. [n = 8 biological replicates (participants) per exercise condition; participants were males aged 26 ± 1.6 yrs and untrained (VO_2_ peak 42.6 ± 1.5 ml/min/kg). Smokers and those participating in regular physical activity (≥ 1 session/week) were excluded.] **(C)** Average percent cysteine modification across subcellular locations in all exercise conditions (pre, post, rec). PM, plasma membrane; nuc, nucleus; mito, mitochondria; golgi, golgi apparatus; ER, endoplasmic reticulum; cyto, cytosol. [n = 24 biological replicates per condition; data are shown as mean.] **(D)** Average percent cysteine modification pre, post, rec in endurance, sprint, and resistance exercise. [n = 8 biological replicates per exercise condition; data are shown as mean ± SEM.] **(E)** Proportion of proteins with 0, 1, 2, ≥3 highly modified cysteines (>10%). **(F)** Heatmap of percent change in cysteine oxidation from respective pre-exercise, post and recovery from all samples (n = 8/group). Insets display percent change in cysteine oxidation of exercise specific regulation in recovery and post exercise. **(G)** Correlation matrix of percent change in individual cysteines from respective pre-exercise across exercise interventions (endurance, sprint, and resistance) and time (post and rec). **(H)** Proportion of proteins that contain ≥1 highly modified cysteine (>10%). **(I-J)** Pairwise comparison of cysteine modification state between sprint post and recovery vs. the pre-exercise proteome. [n = 8 biological replicates per condition. Differentially modified cysteines were identified by two-sided paired Student’s t-test; highlighted/colored cysteines indicate sites reaching |Δ% modification| > 10%.] **(K)** Pairwise comparison of cysteine modification state between endurance post vs. resistance post proteome. [n = 8 biological replicates per condition; statistical significance assessed as in (I-J).] **(L)** Pairwise comparison of cysteine modification state between sprint post vs. endurance post proteome. [n = 8 biological replicates per condition; statistical significance assessed as in (I-J).] Data are presented as mean ± SEM unless otherwise indicated. All comparisons use n = 8 biological replicates (participants) per exercise condition. Statistical significance of cysteine oxidation changes was assessed by two-sided paired Student’s t-test; cysteines passing P < 0.05 are annotated as significantly modified, cysteine passing P < 0.05 and |Δ% modification| > 10% are annotated as highly modified.

In this study, we applied CPT-MS to systematically map the landscape of protein cysteine targets of redox regulation during exercise in human muscle **(Figure 1A)**. This approach was coupled with tandem mass tag (TMT) multiplexing, enabling the simultaneous analysis of eight biological replicates within a single experiment **(Figure 1B; see Methods)**. To profile proteome-wide cysteine oxidation regulated by exercise in human muscle, we subjected eight healthy untrained young men (age, 26.3 ± 1.3 years; BMI, 23.5 ± 0.7 kg/m^2^; maximal oxygen uptake [VO_2_ max], 42.6 ± 1.5 mL/kg/min) to three separate exercise interventions in a randomized crossover design with a 2-week washout between each bout **(Figure 1B & Table S1)**. These interventions included an acute bout of endurance (90 min, 60% VO_2_ max), sprint (3 x 30 s all-out cycling), or resistance exercise (6 sets of 10 repetition maximum knee extensions). Muscle biopsies were obtained from the vastus lateralis before (pre), immediately after (post) and 3 h after exercise cessation (rec) as detailed in Methods. Tissues were processed using the CPT redox proteomics workflow **(Figure 1A)**. The resulting dataset, termed OxiMuscle, included 9,177 unique cysteine sites on 2,782 proteins, quantified across 17,492 individual site measurements spanning all exercise interventions **(Table S2)**.

### Population Characteristics of Redox-Regulated Muscle Proteins Upon Exercise

The overall cellular and compartmental redox tone was remarkably similar between sedentary muscle and all forms of exercise **(Figure 1C)**. Interestingly, no exercise intervention induced bulk protein cysteine oxidation **(Figure 1D)**. Furthermore, at the organelle level, no form of exercise significantly altered the bulk subcellular redox tone in muscle **(Figure S1B)**. We performed protein- and site-level analyses to identify specific populations of proteins that are redox-regulated by different forms of exercise in muscle. In all exercise interventions, most cysteine residues exhibited no exercise-dependent change in redox modification **(Figure 1E)**. However, between 2% and 7% of proteins contained at least one cysteine residue that was highly regulated (greater than 10% change) in each exercise modality. Each exercise intervention drove a sub-population of cysteines to be highly modified, and in the entire dataset, there were 697 exercise-regulated cysteine residues, accounting for 8% of the 9,177 unique cysteine sites mapped **(Figures 1F**; **Figure S1C)**. Interestingly, many of the proteins containing highly modified cysteines regulated by exercise differed depending on the intervention as there was weak correlation of highly modified sites across exercise types **(Figure 1G)**, with sprint exercise exhibiting the highest number of modified cysteine residues **(Figure 1H)**. The extent of cysteine modification was independent of protein abundance in every exercise intervention **(Figure S1D)**.

### Exercise-mediated Redox Regulation of Individual Proteins is Intervention-Specific

While exercise induced only modest bulk changes at the population level in the redox proteome across all interventions **(Figure 1D)**, we observed individual cysteine residues that exhibited substantial modification in at least one form of exercise **(Figure 1I-J; Figure S1E-H)**. A fraction of these exercise-regulated cysteines exhibited a high degree of specificity, with each form of exercise displaying modifications distinct to that specific intervention **(Figure 1F)**. Additionally, when performing pairwise comparisons of exercise interventions post or following recovery, we identified distinct clusters of highly modified cysteine sites induced by different exercise types **(Figure 1K-L; Figure S1I-J)**. We assigned a score to each cysteine residue within the OxiMuscle dataset based on the degree of dynamic oxidation driven by each form of exercise. This analysis showed that of the 697 sites exhibiting a high degree of dynamic and exercise-specific modification, 407 occurred in sprinting exercise, 185 in endurance exercise, and 105 in resistance exercise **(Table S3, Figure S1K)**. These findings provide evidence that upon exercise select protein cysteines are subject to dynamic redox regulation as opposed to broad oxidation of the muscle proteome.

The observation that distinct exercise modalities drive oxidation of different subsets of cysteine residues likely reflects, in part, differences in the subcellular sources and timing of ROS initiation by each form of exercise. Available evidence does not support an increase in mitochondrial ROS during high-intensity burst exercise^11,24^. Instead, NADPH oxidases (NOX), localized to the sarcolemma, transverse tubules, and sarcoplasmic reticulum^13,25^, appear to be the predominant ROS sources in these contexts. In particular, NOX2^11^ and NOX4^13^ are required for acute moderate-intensity exercise-induced cytosolic and whole-muscle ROS production in mice. Notably, mitochondrial ROS increase upon prolonged repeated exercise bouts^26^ and during immediate recovery in the hours following exercise cessation^27^, indicating that subcellular ROS dynamics are both intensity- and time-dependent. These modality-dependent differences in the sources, magnitude, and kinetics of ROS production may provide an explanation for the exercise-specific patterns of cysteine oxidation observed in the OxiMuscle dataset.

### Redox Regulation of Established Muscle Exercise Adaptation Pathways

To assess whether exercise-regulated cysteine oxidation sites converge on coordinated biological activities, we integrated the OxiMuscle dataset with established protein networks (BioPlex 3.0) and Gene Ontology (GO) pathway analysis^28^ **(Figure 2A)**. Protein networks (BioPlex) or pathways (GO) exhibiting coordinated redox regulation of cysteines upon any form of exercise, defined by a shift in oxidation rate exceeding 10%, were classified as “exercise-regulated redox networks” **(Table S4 and Figure 2B).**

**Figure 2.**
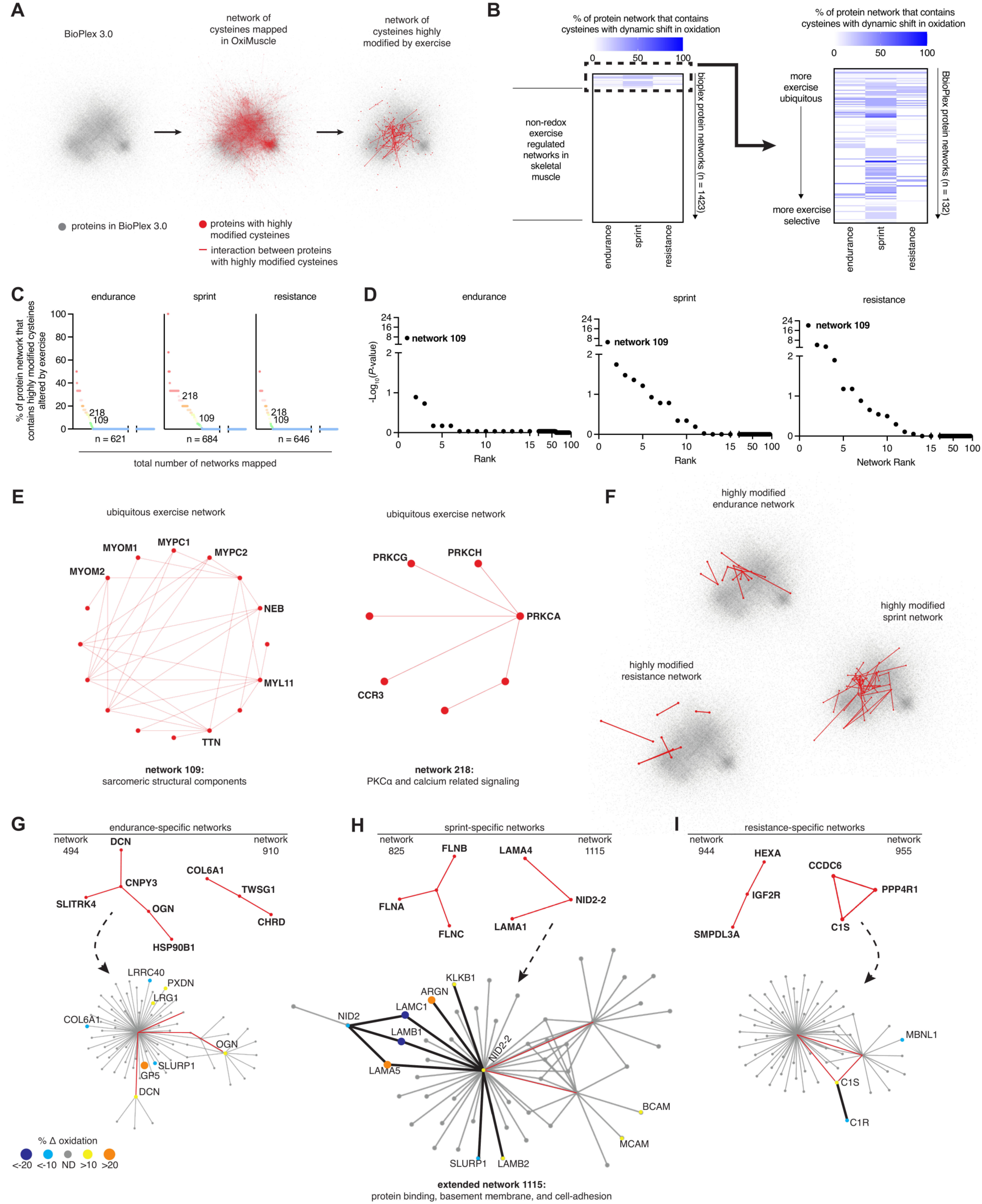
Mapping OxiMuscle to Protein-Protein Interaction Networks Identifies Ubiquitous and Unique Exercise-Influenced Communities. **(A)** Proteins included in the BioPlex 3.0 (HEK and HCT) protein interaction networks (gray dots) overlaid with all proteins found in OxiMuscle (red dots) and known interactions found between proteins that were also identified in OxiMuscle (red lines). OxiMuscle data were further filtered to proteins and interactions with only highly modified cysteines (>10%). **(B)** Left: Percentage of protein networks in BioPlex 3.0 that contain a cysteine that was found highly modified vs. respective pre-exercise in OxiMuscle ranked by coefficient of variation (CV) to determine exercise-selective networks and those not subject to exercise-induced redox modification. Skeletal muscle constitutes a small protein pool compared to the 1423 networks found in BioPlex 3.0 containing many tissue-specific protein interactions. Right: Expanded view of the protein networks in BioPlex 3.0 with a cysteine highly modified vs. respective pre-exercise and ranked by CV. **(C)** Percent of protein networks containing a highly modified cysteine (>10%) in each exercise type identified networks 109 and 218 as the only two found in all three exercise conditions. [n = 8 biological replicates per exercise condition.] **(D)** In each exercise type, all BioPlex 3.0 networks ranked by Fisher’s exact test significance determining if a protein was highly modified (>10%), significantly changed compared to pre-exercise, and was found in each network. Network 109 was the most significant in each exercise type. [n = 8 biological replicates per condition; Fisher’s exact test P-values corrected for multiple testing by Benjamini-Hochberg FDR; significance threshold FDR-adjusted P < 0.05.] **(E)** The two networks containing highly modified cysteines (>10%) in all exercise types, 109 and 218, were found to consist of structural components of sarcomeres and calcium signaling, respectively. **(F)** Exercise-specific networks containing highly modified cysteines (>10%). **(G-I)** Two representative networks specific to each exercise modality and their respective protein-protein interaction branches (extended networks). Black lines indicate a known BioPlex interaction and includes two proteins with a highly modified cysteine (>10%) found in OxiMuscle. All analyses use n = 8 biological replicates (participants) per exercise condition. Network enrichment in **(D)** was assessed by Fisher’s exact test with Benjamini-Hochberg FDR correction; networks with FDR-adjusted P < 0.05 were considered significantly exercise-regulated.

Exercise-regulated redox networks identified across multiple interventions coalesced within a network of proteins constituting structural components involved in muscle contraction and calcium signaling **(Figures 2C-E)**. These included sarcomeric proteins that provide the scaffolding for cross-bridge formation and force transmission during contraction^29–34^. These findings are consistent with prior evidence that ROS regulates the contractile apparatus to tune muscle function. Titin, in particular, contains cysteines within its immunoglobulin domains that undergo S-glutathionylation upon force-induced unfolding, altering passive stiffness and elasticity^35,32,36^, while myosin can be S-glutathionylated *in vitro* with consequences for ATPase activity^32^, and cardiac myosin light chain 1 and cardiac myosin binding protein C are S-glutathionylated at specific cysteines with functional consequences for myofilament calcium sensitivity^37–39^.

Exercise-regulated redox networks also included proteins involved in calcium-dependent and receptor-mediated signaling pathways **(Figure 2E)**, including protein kinase C (PKC) enzymes. PKCs are redox-sensitive kinases involved in calcium handling that contain cysteine-rich C1 domains that coordinate zinc and undergo oxidative modification, disrupting zinc binding and activating kinase activity independently of canonical diacylglycerol signaling^40–44^. In addition to oxidative activation, PKC isoforms can undergo S-glutathionylation, leading to isoform-specific inhibition and highlighting divergent regulatory roles of redox modifications across the PKC family^45^. Calcium release during muscle contraction activates PKCs, which enhance cross-bridge cycling through phosphorylation of contractile proteins such as troponin^39^, suggesting that exercise may simultaneously impose calcium- and redox-dependent modifications to activate PKC signaling during contraction.

Together, our findings show that exercise-induced ROS modify key regulatory proteins across the sarcomere, spanning structural, mechanical, and calcium-regulated proteins central to tissue adaptation during muscle contraction. In addition to protein networks regulated across all exercise types, we identified networks selectively regulated in specific forms of exercise **(Figure 2F)**. An example of a prominent network specific to endurance exercise contained proteins related to extracellular matrix structure and toll-like receptor signaling **(Figure 2G)**. Conversely, networks specific to sprint and resistance exercise included basement membrane and cell-adhesion proteins, and complement pathway proteins, respectively **(Figure 2H, I)**.

In parallel to protein network analysis, we systematically curated cysteine oxidation sites regulated by exercise to identify those that coalesced to distinct biological activities as determined by GO analysis **(Figure 3A)**. These sites were then mapped to 30 pathways known to be responsive to exercise interventions and 8 pathways related to general cell biological processes **(Figure 3A-D; see Methods)**. We observed regulation of cysteines within proteins regulating glycolytic metabolism following sprint exercise **(Figure 3E)** that resolves upon recovery **(Figure S2A)**. Similarly, we observed coordinated modification of oxidative phosphorylation related proteins following endurance exercise **(Figure 3F)**, that renormalized upon recovery **(Figure S2B)**. For resistance exercise, we observed modification of proteins involved in protein synthesis **(Figure 3G)**, an effect which persisted into the recovery phase **(Figure S2C)**. We also observed dynamic changes in extracellular matrix proteins in the responses **(Figure S2D-E)**. The above-described processes are well known to be relevant in the context of muscle adaptation to exercise. These exercise-specific redox signatures coalesce on proteins central to established metabolic adaptations inherent to muscle adaptation to exercise. Specifically, oxidative phosphorylation is a hallmark of endurance adaptation^46–51^, mTORC1-mediated protein synthesis is elevated upon resistance exercise^52,53^, and glycolytic flux is a major driver of ATP production during high-intensity sprinting^51,54,55^. Together, our curation of exercise-regulated cysteine oxidation sites across these pathways provides a resource for interrogating how ROS-dependent signaling intersects with the molecular machinery driving metabolic, structural, and translational adaptations to distinct forms of exercise.

**Figure 3.**
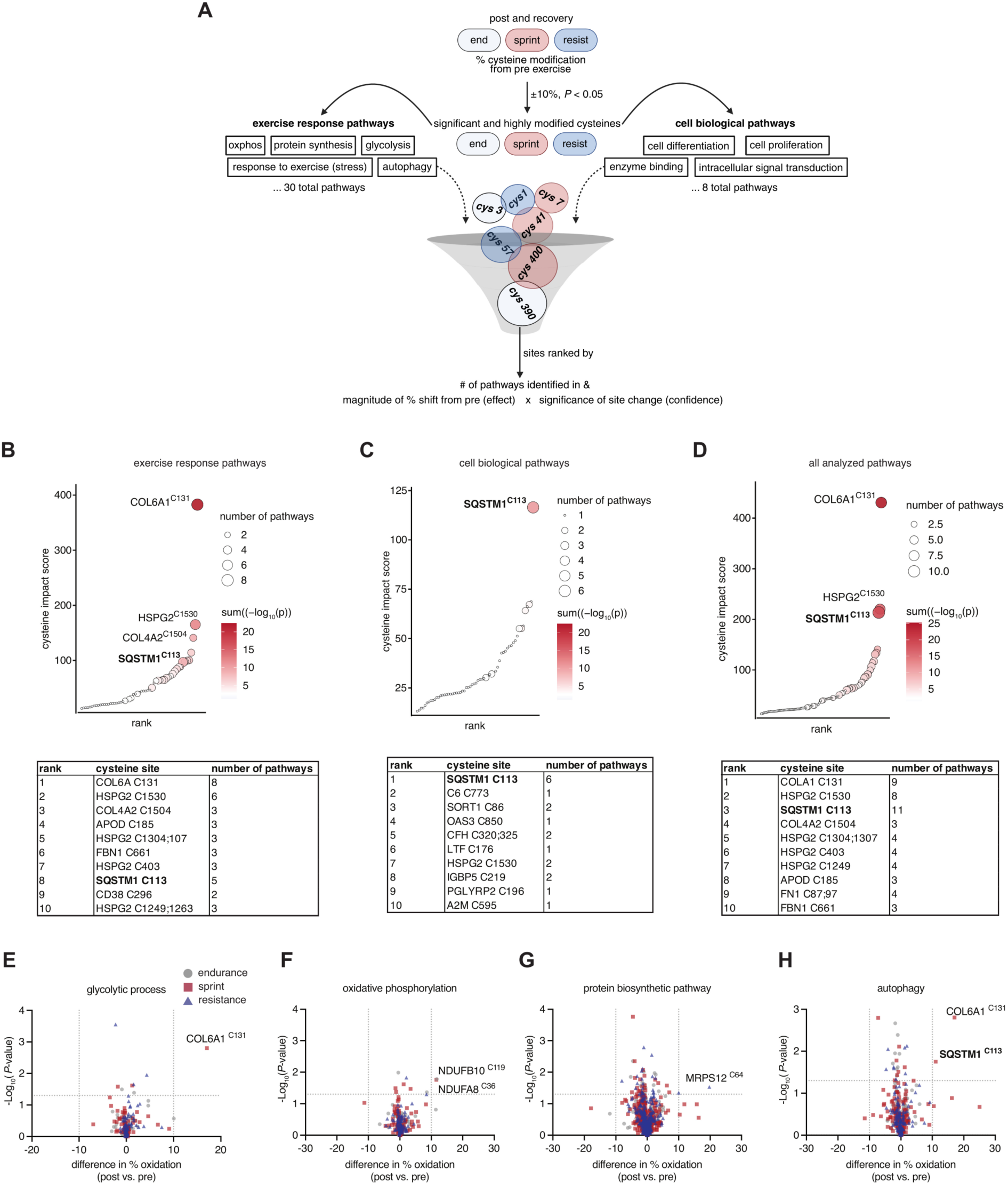
OxiMuscle Confirms Dynamic Modification of Proteins Involved in Exercise Metabolism. **(A)** Diagram of cysteine site filtering through Gene Ontology (GO) pathways to determine potential influential cysteine sites across exercise-adaptive responses and diverse biological processes. **(B-D)** Exercise-response pathways, cell-biological pathways, and all analyzed pathways ranked by cysteine impact score (Sum of absolute % modification shift from pre-exercise multiplied by -log_10_ (P-value) for each pathway in a set a cysteine site appears). **(E-H)** Pairwise comparison of cysteine modification state post vs. the pre-exercise proteome in each exercise type and filtered for proteins in the glycolytic, oxidative phosphorylation, the protein biosynthetic and the autophagy GO process. [n = 8 biological replicates per exercise condition; highlighted cysteines reach P < 0.05 (two-sided paired Student’s t-test) and |Δ% modification| > 10%.] All analyses use n = 8 biological replicates (participants) per exercise condition. For **(B-D),** per-cysteine P-values were derived from two-sided paired Student’s t-tests comparing post- or recovery- to pre-exercise oxidation; impact scores sum absolute Δ% modification × -log_10_ (P-value) across cysteines mapping to each process. For (**E-H)**, highlighted cysteines pass P < 0.05 (two-sided paired Student’s t-tests) and |Δ% modification| > 10%.

### OxiMuscle Uncovers a Redox Switch that Regulates Autophagosome Formation Through p62

We next focused our attention on regulated sites nominated by our systematic analysis contained within proteins likely to be central to the muscle response to exercise. Our above analysis revealed that among the highly redox-modified cysteines induced upon exercise is Cys113 on the central autophagy protein p62 **(Figure 4A)**. SQSTM1/p62 is a prototypic autophagic receptor that links ubiquitylated substrates to nascent autophagic vesicles^56,57^. Interestingly, we observed increased oxidation of p62 immediately following sprinting exercise **(Figure 3H)** that resolved during recovery **(Figure S2F)**. p62 cysteine oxidation appeared specific to Cys113, as other cysteines on p62 did not show significant changes **(Figure 4A)**. This site was of particular interest to us as autophagy is required for exercise training-induced skeletal muscle remodeling and improvement of physical performance^58–61^. Moreover, exercise-induced autophagy is also required for muscle glucose homeostasis^59^ and insulin sensitivity^60^, and training-induced autophagy accompanies mitochondrial biogenesis^58^.

**Figure 4.**
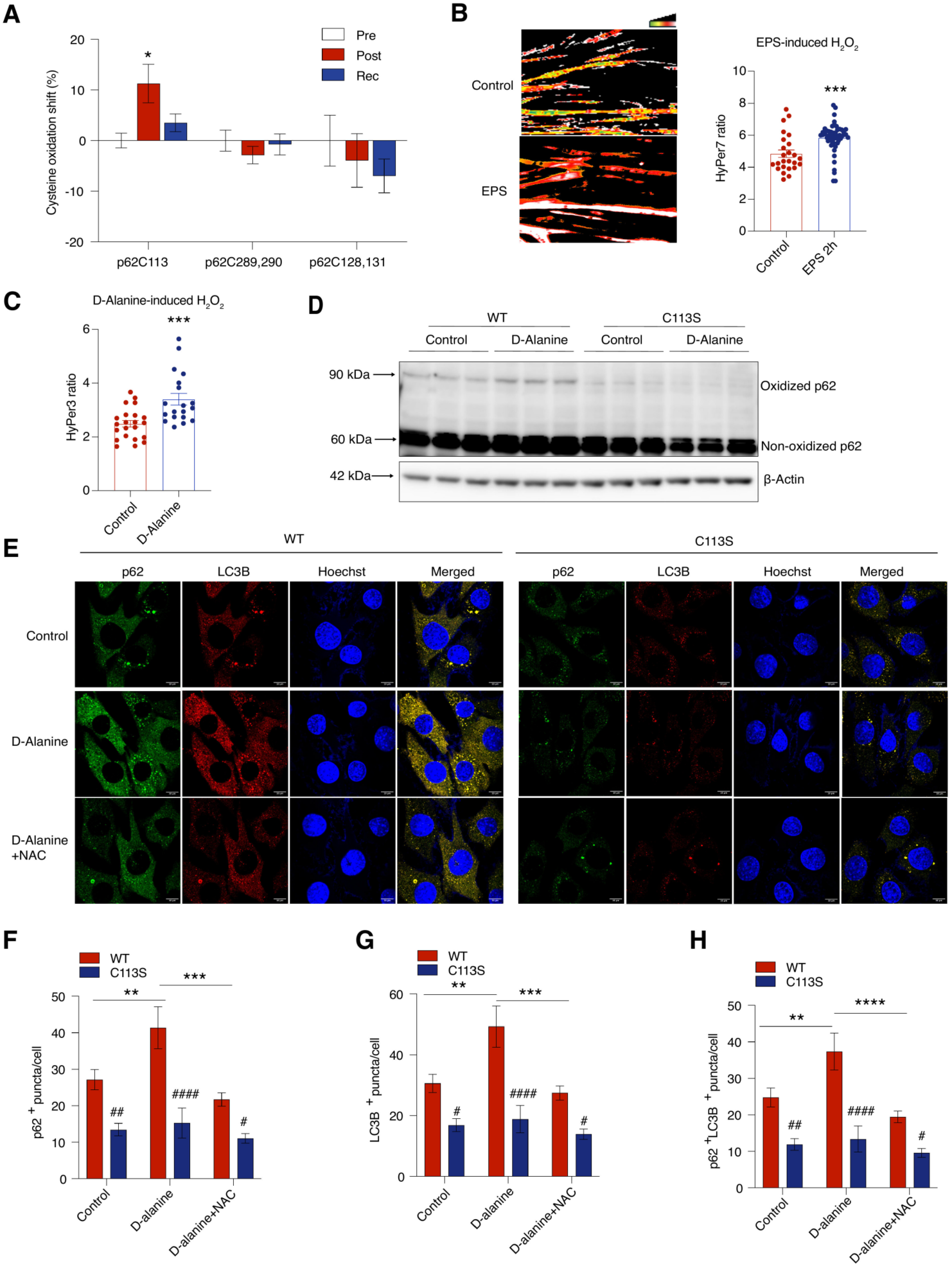
OxiMuscle Uncovers a Redox Switch That Regulates Autophagosome Formation Through p62 Cysteine 113. **(A)** Average percentage shift in cysteine oxidation at p62 Cys113, Cys289/290 and Cys128/131 in human skeletal muscle collected before exercise, immediately after sprint exercise, and 3 hours post-exercise. [n = 8 biological replicates (participants); data are shown as mean ± SEM; statistical significance assessed by one-way repeated-measures ANOVA with Tukey’s post-hoc test.] **(B)** H₂O₂ production in HyPer7-DAAO-transduced myotubes following electrical pulse stimulation (EPS) for 2 hours. [Data shown as mean ± SEM; statistical significance assessed by two-sided Student’s t-test]. **(C)** H₂O₂ production in HyPer3-DAAO-transduced myotubes following D-Alanine treatment for 2 hours. [Data shown as mean ± SEM; statistical significance assessed by two-sided Student’s t-test.] **(D)** Western blot showing oxidized and non-oxidized p62 expression in WT and C113S myoblasts treated with D-Alanine. β-actin was used as loading control. **(E)** Representative confocal images of p62 and LC3B in WT and C113S myoblasts expressing DAAO and treated with D-Alanine and NAC for 2 hours in the presence of chloroquine. [25 fields per experiment, 2-6 cells per field. Scale bar = 10 μm.] **(F-H)** Quantification of p62⁺ puncta, LC3B⁺ puncta, and double-positive (p62⁺LC3B⁺) puncta in WT and C113S myoblasts expressing DAAO treated with D-Alanine and N-acetylcysteine (NAC) for 2 hours in the presence of chloroquine. [Data shown as mean ± SEM; statistical significance assessed by two-way ANOVA with Tukey’s post hoc test (genotype × treatment). *comparison within the same genotype; ^#^comparison between genotypes at the same time point.] For **(A)**, n = 8 biological replicates (participants). For **(B-H)**, n = 3 independent biological replicates (separate differentiations/transductions) with 3 technical replicates per experiment. Statistical significance was assessed by one-way repeated-measures ANOVA with Tukey’s post hoc test **(A)**, two-sided Student’s t-test **(B-C)** or two-way ANOVA with Tukey’s post-hoc test **(F-H)**; *P < 0.05, **P < 0.01, ***P < 0.001, ****P < 0.0001; ^#^P < 0.05, ^##^P < 0.01, ^###^P < 0.001, ^####^P < 0.0001.

The influence of p62 oxidation on exercise-driven muscle adaptation is unknown, and whether contraction-induced cysteine oxidation regulates p62 function remains unexplored. More generally, p62 oxidation-induced oligomerization is known to enhance autophagy and promote cell survival under oxidative stress^62^. The N-terminal region of p62 (amino acids 1–122) can form disulfide-linked conjugates in response to oxidative stress. This region contains five cysteine residues (Cys26, Cys27, Cys44, Cys105 and Cys113), among which Cys105 and Cys113 are the most highly conserved. Cys113 lies in the linker between the PB1 domain (residues 3-102) and the ZZ-type zinc finger (residues 123-173) and is conserved at the equivalent position in mouse p62^62^. Notably, mutation of either Cys105 or Cys113 partially impairs p62 oligomerization driven by exogenously applied oxidants in cells^62^. Polyubiquitin chain-induced p62 filament formation drives autophagic cargo concentration and segregation^63,64^. In addition, the p62 ZZ domain recognizes N-terminally arginylated cargoes, and Cys113 mediates the disulfide-linked p62 polymerization triggered by that binding^65^.

On this basis, we examined whether oxidation of Cys113 regulated p62 function and autophagy in muscle. As a first step, we explored whether oxidation of p62 Cys113 induced by exercise *in vivo* could be recapitulated in cellular models of myotube contractile activity. To do so, we performed acute electrical pulse stimulation (EPS) of C2C12 myotubes. This intervention is known to recapitulate major aspects of muscle adaptations to exercise, including induction of glucose uptake, increased insulin sensitivity and increased sarcomere assembly^66–68^. EPS of C2C12 myotubes was sufficient to increase intracellular ROS^69,70^ **(Figure 4B)**. In parallel, we used a chemogenetic approach to acutely elevate myotube hydrogen peroxide levels. This method relies on ectopic expression of R. *gracilis* D-amino acid oxidase (DAAO), which catalyzes the conversion of D-amino acids to their corresponding alpha-keto acids to produce hydrogen peroxide^69,71^. In DAAO-expressing myotubes, D-alanine drove acute elevation of hydrogen peroxide **(Figure 4C)**. Interestingly, we observed that DAAO-produced hydrogen peroxide drove accumulation of a higher molecular weight form of p62 **(Figure 4D)**. To test whether this ROS-induced form of p62 required Cys113, we engineered C2C12 cells for which p62 Cys113 was replaced with serine (C113S), rendering this site recalcitrant to redox modification **(Figure S3A)**. Mutation of Cys113 resulted in loss of this higher MW form of p62 **(Figure 4D)**. We next tested whether oxidation of p62 Cys113 played a role in autophagosome formation and autophagy in myoblasts and myotubes. p62 mediates binding of ubiquitinated proteins to nascent autophagic membranes to promote formation of the autophagosome around this cargo^56,72^. Tracking endogenous p62 and LC3B, a marker of autophagic membranes^73,74^, we found that both basal and intracellular ROS-induced p62 and LC3B puncta were significantly diminished in p62 C113S myoblasts **(Figure 4E-H)**. We next examined whether pharmacologic manipulation of thiol redox state induced similar effects. We found that the thiol reducing agent N-acetyl cysteine (NAC) inhibited accumulation of p62 puncta, LC3B and double positive p62^+^LC3B^+^ puncta **(Figure 4E-H)**. Notably, there was no significant additive inhibitory effect of NAC in C113S cells, suggesting that the thiol reducing effects on these parameters occurred via regulation of p62 Cys113.

In parallel, we used the Halo-GFP-LC3 dual fluorescence system to track autophagosome formation and autophagic flux^75^. In myoblasts, autophagosome number (GFP^+^ Halo^+^ puncta) was substantially lower in p62 C113S cells. Conversely, autophagic flux in terms of the autophagosome number entering the lysosome (Halo^+^GFP^−^ puncta) was comparable **(Figure 5A-B)**. In myotubes, basal autophagosome number and autophagic flux were substantially lower in C113S cells **(Figure 5C-D).** Moreover, EPS induced autophagosome biogenesis in WT myotubes, and this effect was significantly inhibited in p62 C113S myotubes **(Figure 5C-D)**. Upon EPS, the autophagic flux ratio in C113S myotubes was significantly inhibited compared to WT **(Figure 5C-D)**. Taken together, these data demonstrate that oxidation of p62 Cys113 plays a critical role in autophagosome formation and induction of autophagy upon myotube contraction.

**Figure 5.**
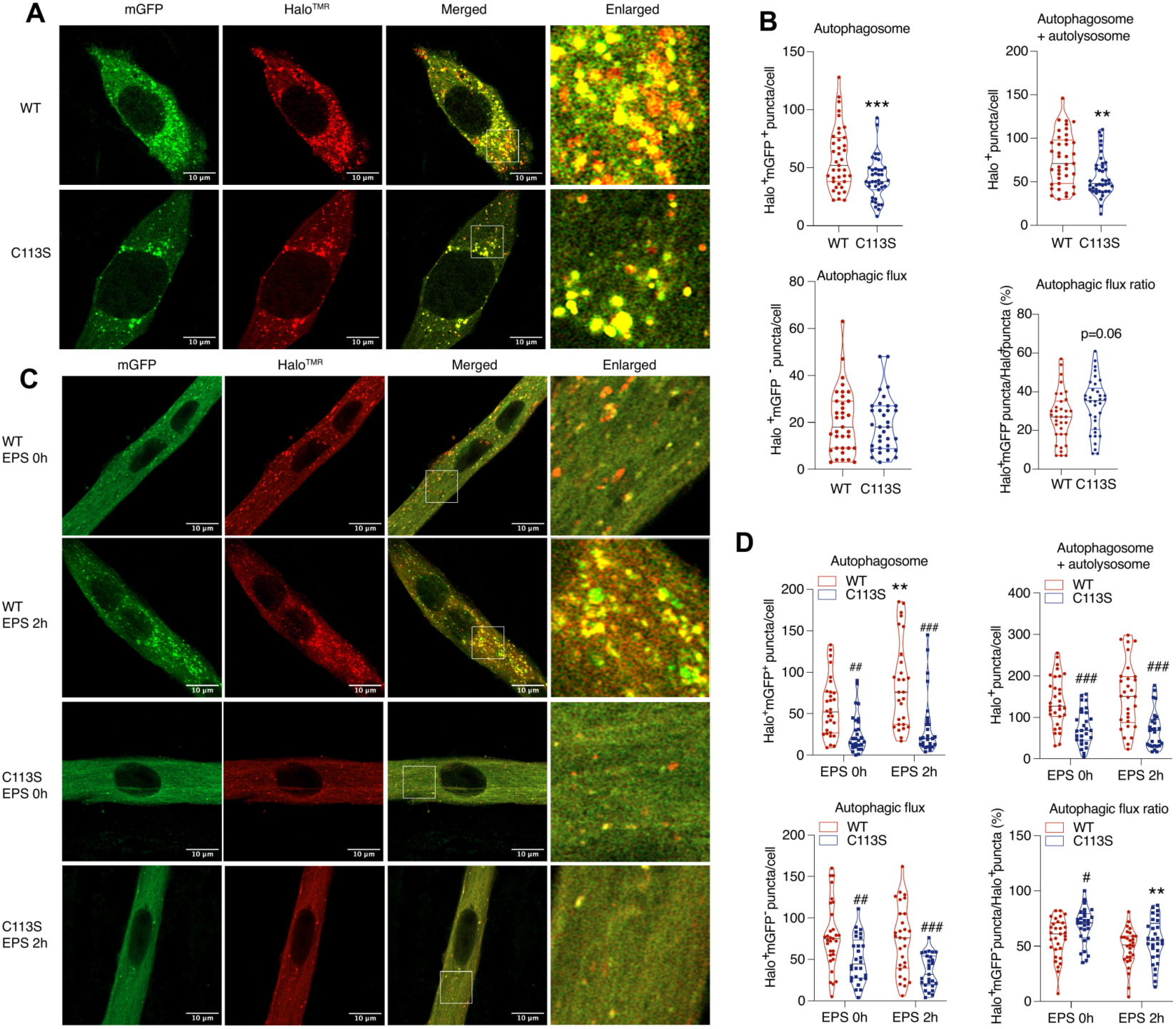
p62 Cys 113 is Required for Basal and H_2_O_2_-Induced Autophagosome Formation. **(A)** Representative confocal images of mGFP and Halo^TMR^ in WT and C113S myoblasts expressing HaloTag7-mGFP-LC3 and labeled with TMR-conjugated Halo ligand. Scale bar = 10 μm. **(B)** Quantification of double-positive (Halo⁺mGFP⁺) puncta, total Halo⁺ puncta, and Halo⁺mGFP⁻ puncta in WT and C113S myoblasts expressing HaloTag7-mGFP-LC3 and labeled with TMR-conjugated Halo ligand. [n = 40–44 cells per genotype; data shown as mean ± SEM; each dot represents a single cell/field; statistical significance assessed by two-sided Student’s t-test.] **(C)** Representative confocal images of mGFP and HaloTMR in WT and C113S differentiated myotubes expressing HaloTag7-mGFP-LC3, labeled with TMR-conjugated Halo ligand, and subjected to electrical pulse stimulation (EPS) for 2 hours. Scale bar = 10 μm. **(D)** Quantification of double-positive (Halo⁺mGFP⁺) puncta, total Halo⁺ puncta, and Halo⁺mGFP⁻ puncta in differentiated WT and C113S myotubes expressing HaloTag7-mGFP-LC3, labeled with TMR-conjugated Halo ligand, and subjected to EPS for 2 hours. [n = 30 myotubes per genotype/condition; data shown as mean ± SEM; each dot represents a single myotube; statistical significance assessed by two-way ANOVA with Tukey’s post-hoc test (genotype × EPS). *comparison within the same genotype; ^#^comparison between genotypes at the same time point.] n = 3 independent biological replicates (separate differentiations/transfection) per genotype/condition. Statistical significance was assessed by two-sided Student’s t-test **(B)** or two-way ANOVA with Tukey’s post-hoc test **(D)**; *P < 0.05, **P < 0.01, ***P < 0.001; ^#^P < 0.05, ^##^P < 0.01, ^###^P < 0.001.

### p62 Cys113 Regulates Muscle Adaptation to Exercise *In Vivo*

To test whether oxidation of p62 Cys113 is required for muscle adaptation to exercise *in vivo*, we generated a mouse line in which the endogenous *Sqstm1* cysteine at position 113 is replaced with serine (p62 C113S). Correctly targeted founders were backcrossed to C57BL/6J, and heterozygous offspring were intercrossed to generate homozygous p62 C113S and littermate wild-type (WT) animals used in all subsequent experiments.

We verified the C113S substitution by Sanger sequencing of the *Sqstm1* locus from genomic DNA (**Figure 6A**). Intercrosses of p62 C113S heterozygous mice yielded viable homozygous offspring, although homozygous animals were recovered at lower-than-expected Mendelian frequency (**Table S5**), suggesting that loss of p62 Cys113 may partially impair developmental or early postnatal fitness. p62 C113S mice were born healthy, were grossly indistinguishable from WT littermates, and displayed no overt physical or behavioral abnormalities. Steady state p62 protein abundance in skeletal muscle (gastrocnemius and soleus) was comparable between genotypes by proteomics (**Figure 6B**) and immunoblot (**Figure 6C**).

**Figure 6.**
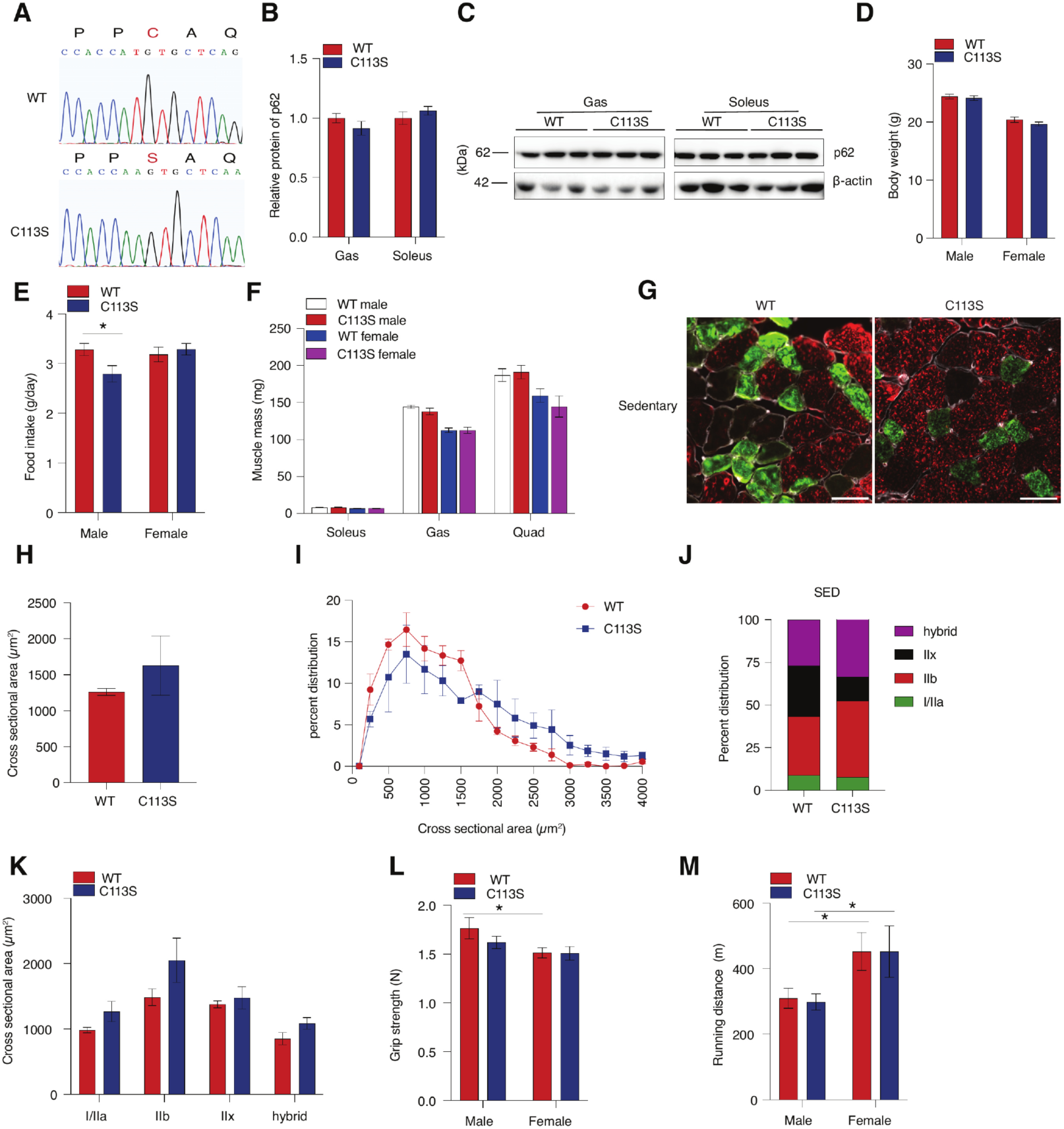
Characterization of sedentary WT and p62 C113S mice. **(A)** Representative Sanger sequencing chromatograms confirming the C113S substitution in homozygous p62 C113S mice. **(B)** Quantification of p62 protein abundance in gastrocnemius (Gas) and soleus muscles from WT and C113S mice by proteomics. [n = 9–11 mice per group.] **(C)** Representative immunoblot and quantification of p62 protein abundance in gastrocnemius and soleus muscle lysates from WT and C113S mice. β-actin was used as loading control. [n = 3 mice per group.] **(D)** Body weight (g) of 10–12-week-old male and female WT and C113S mice. [n = 9–11 per group.] **(E)** Daily food intake (g/day) in male and female WT and C113S mice. [n = 9–11 per group.] **(F)** Wet muscle mass (mg) of Soleus, gastrocnemius (Gas), and quadriceps (Quad) in male and female WT and C113S mice. [n = 9–11 per group.] **(G)** Representative fiber-type immunofluorescence images of plantaris cross-sections from sedentary WT and C113S mice. [n = 3 mice per group.] **(H)** Mean fiber cross-sectional area (CSA, µm²) in plantaris from sedentary WT and C113S mice. [n = 3 mice per group.] **(I)** Frequency distribution of fiber CSA in plantaris from sedentary WT and C113S mice. [n = 3 mice per group.] **(J)** Percent distribution of fiber types (I/IIa, IIb, IIx, hybrid) in sedentary (SED) plantaris from WT and C113S mice. [n = 3 mice per group.] **(K)** Mean fiber CSA (µm²) by fiber type (I/IIa, IIb, IIx, hybrid) in sedentary plantaris from WT and C113S mice. [n = 3 mice per group.] **(L)** Grip strength (N) in male and female WT and C113S mice. [n = 9–11 per group.] **(M)** Running capacity in a treadmill exhaustion test (m) in male and female WT and C113S mice. [n = 9–11 per group.] Data are presented as mean ± SEM unless otherwise indicated. Statistical significance was assessed by two-sided Student’s t-test **(H and J)** or two-way ANOVA with Tukey’s post-hoc test **(B, D-F, K-M)**; *P < 0.05.

By 12 weeks of age, body weight (**Figure 6D**), lean mass (**Figure S3B**) and fat mass (**Figure S3C**) were comparable across genotypes. Food intake (**Figure 6E**), and locomotion distance in cage (**Figure S3D**) were indistinguishable across genotypes in female mice, with male C113S mice showing a trend towards lower food intake and locomotion distance **(Figure 6E and S3D)**. In addition, oxygen consumption (**Figure S3E)**, carbon dioxide production (**Figure S3F)**, respiratory exchange ratio **(Figure S3G)**, and energy expenditure (**Figure S3H**) were comparable in female but lower in male C113S mice. Anatomically, skeletal muscle from sedentary p62 C113S and WT mice was indistinguishable. Masses of hindlimb muscles were comparable between genotypes (**Figure 6F**). Cross-sectional fiber area (CSA) of plantaris was not significantly different between genotypes (**Figure 6G-I**). Moreover, there was no significant difference in plantaris fiber-type distribution or CSA of each type of fiber between genotypes **(Figure 6J-K)**. Baseline grip strength (**Figure 6L**) and running capacity in a treadmill exhaustion test (**Figure 6M**) were likewise indistinguishable between genotypes.

Our *in vitro* data established that Cys113 oxidation is key for contraction-induced p62 recruitment and autophagosome formation: the site is oxidized acutely upon stimulation and reversed in recovery, and blocking its modification, genetically in C113S cells or pharmacologically with NAC, blunts autophagy. Because p62-dependent autophagy is a route by which cells adaptively remodel the proteome, we reasoned that this redox switch could shape muscle proteome content specifically upon exercise training. To test whether p62 Cys113 redox regulation shapes the muscle proteome during exercise *in vivo*, we performed label-free quantitative proteomics on gastrocnemius from p62 C113S and littermate WT mice under sedentary conditions and following 5-week voluntary wheel running exercise (VWR), and compared proteins significantly induced in each genotype compared to sedentary controls.

### Mitochondrial Proteome Remodeling is Impaired in Exercised p62 C113S Muscle

p62 C113S and WT mice ran similar distance (**Figure S3I**) but exhibited distinct proteomic responses to exercise (**Figure 7A)**. We first focused our attention to proteins that accumulated selectively in p62 C113S muscle following exercise, which may be indicative of compromised p62-mediated turnover or alternatively aberrant regulation in the absence of effective induction of autophagy. Of the 141 proteins elevated in C113S but not WT muscle after exercise, 52 (∼37%) are annotated as mitochondrial components based on MitoCarta 3.0 (**Figure S3J**). These mitochondrial proteins spanned subunits of all OXPHOS complexes (**Figure 7B**), mitoribosomal subunits (**Figure 7C**), mitochondrial biogenesis and assembly factors (**Figure 7D**), fatty acid oxidation enzymes (**Figure 7E**), TCA cycle enzymes (**Figure 7F**) and mitochondrial carriers (**Figure 7G**). Nuclear- and mtDNA-encoded proteins were represented within this set, arguing against a selective import or assembly defect. A selective remodeling of mitochondrial protein abundance dependent on p62 Cys113 is consistent with prior evidence that exercise induces mitophagy in skeletal muscle^27,59,76,77^ and with the established capacity of mitochondria to undergo piecemeal, cargo-selective turnover via mitochondria-derived vesicles and preferential respiratory chain degradation^78–81^. In addition, upon loss of p62 Cys113, multiple sarcomeric proteins accumulated **(Figure S3K)** supporting a broader role for p62 Cys113-dependent turnover that extends beyond mitochondria^82^.

**Figure 7.**
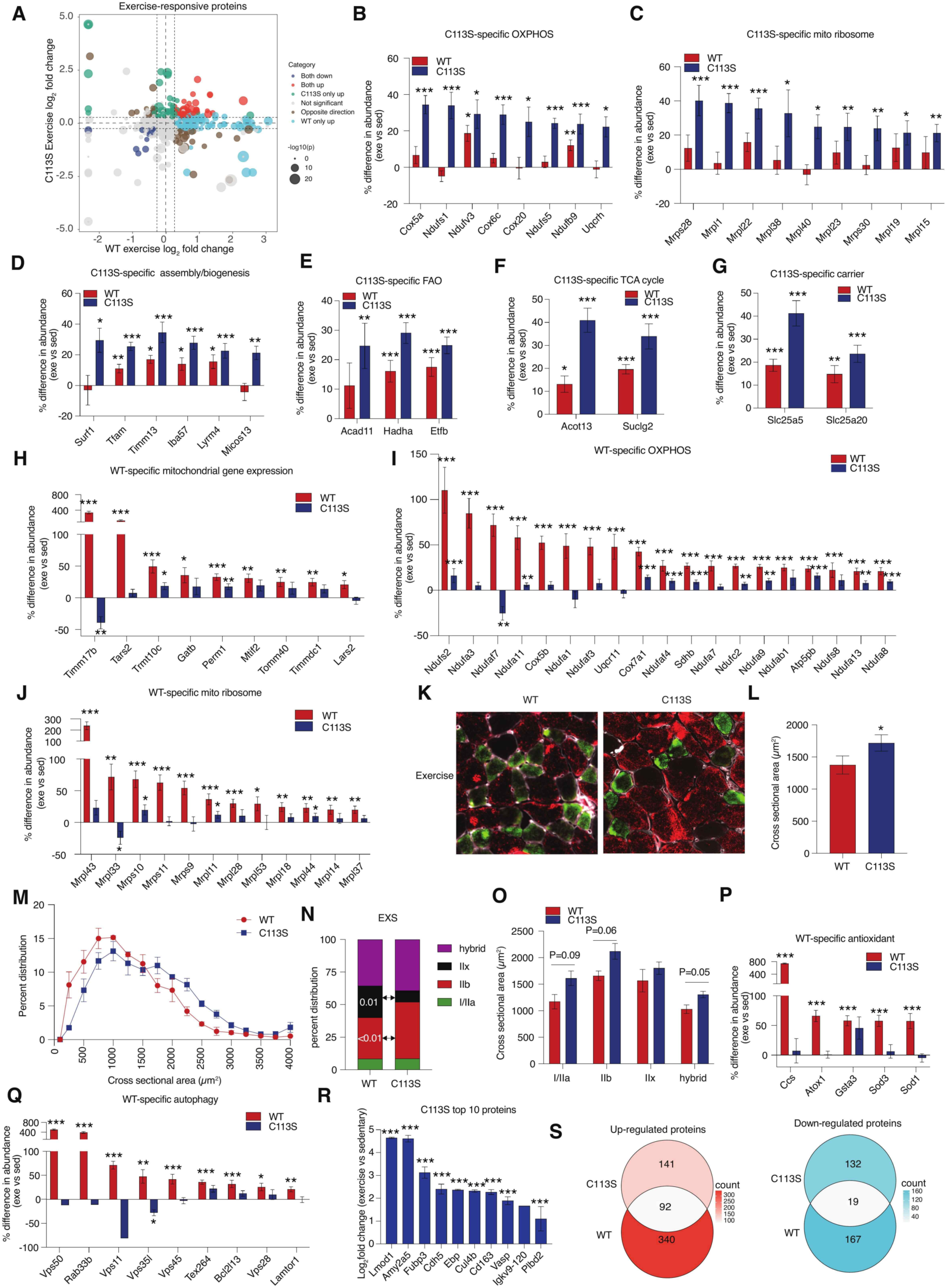
Distinct Mitochondrial and Autophagy Remodeling Responses to Exercise in WT and p62 C113S Skeletal Muscle. **(A)** Scatter plot comparing exercise-induced proteomic changes between WT and p62 C113S skeletal muscle. Proteins were categorized as commonly upregulated, commonly downregulated, WT-specific upregulated, C113S-specific upregulated, oppositely regulated, or not significant. [n = 9–11 per group.] **(B–G)** Percent difference in abundance of C113S-specific mitochondrial pathways in response to exercise, including oxidative phosphorylation (OXPHOS), mitochondrial ribosome, mitochondrial assembly/biogenesis, fatty acid oxidation (FAO), tricarboxylic acid (TCA) cycle, and mitochondrial carrier proteins. [n = 9–11 per group.] **(H)** Percent difference in abundance of WT-specific mitochondrial translation and import machinery. [n = 9–11 per group.] **(I)** Percent difference in abundance of WT-specific OXPHOS subunits. [n = 9–11 per group.] **(J)** Percent difference in abundance of WT-specific mitoribosomal subunits. [n = 9–11 per group.] **(K)** Representative fiber-type immunofluorescence images of plantaris cross-sections from exercised WT and C113S mice. [n = 3 mice per group.] **(L)** Mean fiber CSA (µm²) in plantaris from exercised WT and C113S mice. [n = 3 mice per group.] **(M)** Frequency distribution of fiber CSA in plantaris from exercised WT and C113S mice. [n = 3 mice per group.] **(N)** Percent distribution of fiber types (I/IIa, IIb, IIx, hybrid) in exercised (EXS) plantaris from WT and C113S mice. [n = 3 mice per group.] **(O)** Mean fiber CSA (µm²) by fiber type (I/IIa, IIb, IIx, hybrid) in exercised plantaris from WT and C113S mice. [n = 3 mice per group.] **(P–Q)** Percent difference in abundance of WT-specific antioxidant proteins and autophagy-related proteins. [n = 9–11 per group.] **(R)** Top 10 proteins specifically upregulated in p62 C113S skeletal muscle following exercise. [n = 9–11 per group, data are presented as log_2_ fold change relative to sedentary controls.] **(S)** Venn diagram analysis showing overlap of upregulated and downregulated proteins between WT and p62 C113S mice after exercise. [n = 9–11 per group.] For **(B–J, P and Q)**, bars show mean percent difference in abundance (exercise vs. sedentary) ± SEM per genotype. Statistical significance for each protein was determined by a two-sample Welch’s t-test comparing individual exercised versus sedentary mice within each genotype; For **(L, N, O and R)**, data are presented as mean ± SEM. Statistical significance was assessed by two-sided Student’s t-test; *P < 0.05, **P < 0.01, ***P < 0.001.

Conversely, of the 340 proteins elevated in WT but not C113S muscle after exercise, 132 (∼38%) again map to mitochondria (**Figure S3L**). This set of factors were distinct from the mitochondrial proteins described above and included core components of a mitochondrial biogenic program, such as the mitochondrial gene expression machinery, encompassing RNA processing factors and translation regulators, import/chaperone machinery, and the muscle-restricted PGC-1α/ERR coactivator Perm1 **(Figure 7H)**^83^. Interestingly, distinct components of the oxidative phosphorylation (**Figure 7I**) and mitochondrial ribosome machinery (**Figure 7J**) were selectively increased in WT, which again suggests selective regulation of mitochondrial proteome that depends on p62 Cys113. Together, these data suggest that in WT exercising muscle, mitochondrial proteome remodeling, presumably through a combination of piecemeal mitochondrial clearance and replacement synthesis, is regulated by p62 Cys113.

Interestingly, the altered proteomic remodeling in p62 C113S mice coincided with distinct morphological and myofibrillar features following exercise. Cross-sectional fiber area of plantaris in C113S mice after exercise was larger compared to WT mice (**Figure 7K-M**). Analysis of fiber-type distribution indicated that C113S mice have more fast-type Type IIb and fewer Type IIx fibers, without significant change in slow-type fibers (type I and Type IIa) compared to WT mice (**Figure 7N**). This shift toward a faster, more glycolytic fiber-type profile is consistent with the impaired mitochondrial proteome remodeling described above. Because oxidative fiber identity is tightly coupled to mitochondrial content and biogenic capacity^49,84^, a muscle unable to mount the mitochondrial adaptations that normally accompany training would be expected to default toward a more glycolytic program rather than acquiring or maintaining oxidative character. The fiber-type distribution in exercised C113S muscle thus mirrors, at the morphological level, an apparent compromise in mitochondrial biogenic remodeling evident in the proteomic data. It is noteworthy that the CSA of each type of fiber in C113S mice tended to be greater than in WT mice as well (**Figure 7O**).

### Induction of Autophagy Machinery is Blunted in Exercised p62 C113S Muscle Coincident with Elevation of Inflammatory Markers

Proteins selectively accumulated in exercised WT muscle also included core autophagy factors that fail to rise in exercised p62 C113S muscle^85–87^ (**Figure 7Q**). In addition, a partial Nrf2/antioxidant signature is also evident in WT and muted in C113S (**Figure 7P**); this is consistent with the established role of p62 in KEAP1 sequestration and Nrf2 activation^88–90^. Conversely, among top p62 C113S-specific elevated proteins post exercise were Cd163 and Cdh5/VE-cadherin **(Figure 7R)**, which mark tissue macrophages and vascular endothelium, respectively, rather than myofibers^91,92^. Cd163 is the high-affinity haemoglobin–haptoglobin scavenger receptor of resident and alternatively activated macrophages^93,94^, while Cdh5 is the defining adherens-junction cadherin of the vascular endothelium^95,96^. Their coordinated induction in exercised p62 C113S muscle is likely a consequence of expansion or activation of the macrophage and endothelial compartments, a well-documented consequence of exercise-induced muscle damage and, more acutely, suggestive of disruption of the muscle autophagy-lysosome axis. These changes could therefore reflect downstream readouts of the underlying clearance defect and secondary tissue-level responses to accumulated damaged substrates.

Together, these data suggest that exercise-induced oxidation of p62 Cys113 regulates abundance of mitochondrial, sarcomeric, and aggregate substrates, coupled to the biogenic, antioxidant, and autophagy-machinery programs that remodel the muscle post exercise. Loss of this cysteine does not abolish exercise responsiveness as both genotypes mount substantial and overlapping proteomic remodeling **(Figure 7S)**. In contrast, p62 Cys113 yields a muscle that fails to remodel the abundance of its mitochondrial and sarcomeric proteome compared to WT, perhaps reflecting an uncoupling of mitochondrial clearance from biogenesis, accumulation of structurally and metabolically diverse substrates, and a shift toward secondary inflammatory and vascular signatures rather than adaptive proteostatic remodeling.

## DISCUSSION

In this study we established a quantitative, site-resolved atlas of the human skeletal muscle redox proteome and used it to define the cysteine oxidation events that accompany distinct forms of exercise. The OxiMuscle resource extends prior efforts to catalogue exercise-regulated signaling, principally the phosphoproteome^52,97,98^, into the redox dimension. By quantifying the stoichiometry of reversible oxidation at 9,177 unique cysteine sites on 2,782 proteins, across 17,492 individual site measurements, OxiMuscle moves the field from the premise that exercise generates ROS toward a mechanistic account of which residues those ROS engage.

Three features of the landscape are notable. First, exercise did not produce a bulk oxidative shift of the muscle proteome; instead, most cysteines were unchanged, and regulation was concentrated in a small, high-amplitude subset. This argues against a model of indiscriminate oxidative stress and in favor of targeted, localized redox signaling, consistent with the concept of ROS as spatially confined second messengers rather than diffuse damaging agents^1,2^. Second, some regulated cysteines were substantially intervention-specific, with sprint, endurance, and resistance exercise each engaging distinct residues. This specificity plausibly reflects the differing subcellular sources, magnitude, and kinetics of ROS production across exercise modalities: NADPH oxidases at the sarcolemma, transverse tubules and sarcoplasmic reticulum during high-intensity contraction^11,13,25^ versus mitochondrial sources that rise with prolonged effort and into recovery^26,27^. Third, the regulated sites converged on protein networks and pathways with established roles in the corresponding adaptive response: glycolytic enzymes after sprint; oxidative phosphorylation after endurance; and the protein-synthetic machinery after resistance exercise, indicating that exercise-induced oxidation is not stochastic but is coupled to the metabolic program each stimulus recruits.

Against this backdrop, Cys113 of the autophagy receptor p62/SQSTM1 emerged as a notable node of redox regulation. We show that this residue is acutely and reversibly oxidized by exercise *in vivo*, that its oxidation is required for the assembly of a higher-order p62 species and regulation of autophagosome formation in muscle cells. These findings connect a defined redox event to a physiological output, and they build on prior evidence that the N-terminal region of p62 forms disulfide-linked oligomers under oxidative stress in cells^62^. Our data extend *in vitro* framework in two respects: they demonstrate that this chemistry operates under the physiological ROS fluxes generated by contraction, and they place it within the broader logic of p62 function. Because p62 self-association through its PB1 domain is central to cargo sequestration and autophagosome nucleation^63,64,99^, redox-dependent oligomerization at Cys113 offers an attractive mechanism by which the oxidative tone of contracting muscle could tune the threshold for selective autophagy.

The *in vivo* consequences of disabling this switch were most evident in the mitochondrial proteome. Loss of p62 Cys113 uncoupled two normally coordinated arms of the exercise response: in WT muscle, exercise elevated a coherent mitochondrial biogenic program. In contrast, in p62 C113S muscle a distinct and largely non-overlapping set of mitochondrial proteins accumulated. We interpret this pattern as a failure to couple mitochondrial clearance to replacement synthesis. Exercise is an established stimulus for mitophagy and for piecemeal, cargo-selective removal of mitochondrial constituents via mitochondria-derived vesicles and preferential respiratory-chain turnover, mechanisms established in cultured cells and Drosophila rather than in exercising muscle^27,76–79,81,100^ and the coordination of this turnover with PGC-1α-driven biogenesis is thought to underlie the net mitochondrial remodeling that defines aerobic adaptation. In this light, the p62 Cys113 redox switch is a candidate mechanism linking the acute oxidative signal of contraction to the longer-term maintenance of mitochondrial quality during training.

Distinct mitochondrial proteomic remodeling coincided with muscle morphological differences between genotypes. Following training, p62 C113S plantaris showed a shift toward the fastest, most glycolytic fiber type and a trend toward larger fiber cross-sectional area, whereas sedentary muscle was indistinguishable from WT in fiber size and type. The direction of the fiber-type shift is consistent with a muscle that, when unable to execute oxidative/mitochondrial remodeling upon exercise, defaults toward a more glycolytic program. The larger cross-sectional area should be interpreted with caution and not conflated with beneficial hypertrophy. Impaired autophagy is more commonly associated with the accumulation of damaged proteins and organelles, myofiber dysfunction, and ultimately atrophy than with functional growth^101,102^, and acute inhibition of autophagy does not impair exercise performance or AMPK activation, though autophagy is required to preserve mitochondrial function during damaging contraction^103^. A plausible interpretation is therefore that increased fiber area in C113S muscle reflects the retention of substrates that would normally be cleared, swelling of the proteostatic and organellar content of the fiber, rather than a coordinated anabolic response.

Together, these findings define a chemically specific, physiologically engaged redox switch on p62 that couples the acute oxidative signal of muscle contraction to the selective-autophagy programs that remodel the muscle proteome during training. More broadly, OxiMuscle provides a systematic, quantitative foundation for dissecting how redox signaling is wired into human muscle physiology, and it nominates a catalogue of exercise-regulated cysteines whose functional interrogation may reveal further nodes at which the benefits of exercise are encoded.

### Limitations of Study

Several limitations qualify our above conclusions. The human redox landscape was defined in a cohort of eight young, untrained men, and the generalizability of specific sites to women, older individuals, and trained states remains to be established. The site-occupancy measurements report reversible oxidation in aggregate and do not resolve the specific oxidative modification (sulfenylation, disulfide, S-glutathionylation) at each cysteine. The causal chain from p62 Cys113 oxidation to autophagosome formation, while supported by convergent genetic and pharmacological evidence, is inferred, and the pharmacological reducing agent NAC has thiol-independent actions that cannot be fully excluded. Finally, the *in vivo* phenotyping relied on a single training paradigm and modest group sizes for the morphological analyses; the lower-than-expected Mendelian recovery of homozygotes and the sex-dependent metabolic differences at baseline also indicate roles for p62 Cys113 beyond the exercised muscle that our study does not resolve.

## MATERIALS AND METHODS

### Human exercise protocols

All subjects were fasted overnight prior to any intervention and were familiarized with the three exercise modalities (endurance, sprint, and resistance) prior to testing. During familiarization subjects’ VO_2_ peak and 10 repetition maximum (RM) were determined. All subjects underwent identical exercise protocols in random order, separated by at least 10 days of rest (no exercise). Acute endurance exercise consisted of 90 min of continuous cycling at 60% of VO_2_ peak, sprint exercise consisted of a 5-min warmup at 50W, followed by three bouts of 30-second all-out sprint (Wingate test; workload corresponding to 0.075 kg/kg body mass) on a cycler ergometer, and resistance consisted of a warmup of 3 sets of 10 repetitions at 50% of 10-RM followed by 6 sets of 10 repetitions at a load corresponding to 10-RM with 2 mins of rest between each set. Muscle biopsies, described below, were taken prior to- (pre), immediately post- (post), and following 3h of recovery (rec) for each exercise intervention. Further details of all exercise protocols and trial procedures can be found in Blazev et al^52^.

### Human muscle biopsy and % cysteine oxidation sample preparation

Muscle biopsies were obtained from the vastus lateralis using a 5 mm Bergstrom needle with suction and extracted through separate incisions 5-6 cm apart. 1% lidocaine (AstraZeneca A/S) was administered for local anesthesia in the subcutaneous tissue and fascia before sampling. Muscle biopsies were rapidly flushed in ice-cold saline then snap-frozen in liquid nitrogen and stored at -80°C. Muscle samples (n=8 for each timepoint and exercise type) were homogenized using bead mill in ice-cold 20% trichloroacetic acid (TCA). Each sample was split into equal volume replicates containing ∼200 µg protein and washed sequentially with 20%, 10%, then 5% TCA twice each. One sample replicate was resuspended in 100 mM HEPES, pH 8.5, 2% SDS, 1 mM EDTA, 1 mM DTPA, 10 µM neocuproine, and 35 mM iodoacetamide (IAM), which was eventually used as the oxidized channel (**Figure 1A**). The other sample replicate was resuspended in the same buffer, replacing 35 mM IAM with 35 mM CPT, which was eventually used as the total cysteine channel (**Figure 1A**). Next, both sample replicates were methanol-chloroform precipitated, resuspended in 6.25 mM TCEP for 30 min at 37°C, then treated with 175 mM CPT for 1.5 h at 37°C. Samples were then digested with trypsin and LysC overnight at 37°C with shaking. The samples were then TMT-labeled and processed as described in detail in Xiao et al.^15^. Briefly, samples were desalted, treated with lambda phosphatase, and enriched using immobilized metal affinity chromatography (IMAC). Enriched cysteine-containing peptides were then fractionated into 12 fractions using a high-pH reversed-phase peptide fractionation kit (Thermo Fisher), then combined into 6 fractions (fraction 1 with 7, 2 with 8, 3 with 9, 4 with 10, 5 with 11, and 6 with 12). Lastly, samples were purified via C18 Stage Tip.

### CPT synthesis

CPT synthesis commences with the addition of 6-aminohexylphosphonic acid hydrochloride salt (6-AHP; purchased from SiKÉMIA) to N-succinimidyl iodoacetate (SIA; purchased from Combi-Blocks) to final concentrations of 175 mM 6-AHP and 45 mM SIA. 6-AHP is dissolved in 50 mM triethylammonium bicarbonate (TEAB) and SIA in dimethyl sulfoxide (DMSO) prior to mixing. The reaction is allowed to occur for 1 h at room temperature, protected from light with gentle shaking. The reaction is quenched using 0.1-1% formic acid (FA) added until pH reaches 3-3.5, and the final concentration of DMSO is below 5%. CPT was purified on a 10 g C18 Sep-Pak column (Waters). Briefly, the column was equilibrated with 100% methanol, conditioned with 0.1% FA, followed by CPT addition, washed with 0.1% FA/5% acetonitrile (ACN), and finally eluted with 0.1% FA/50% ACN. CPT was lyophilized at 4°C. An extensive analysis of CPT performance can be found in Xiao et al.,^15^.

### Liquid chromatography and mass spectrometry

Proteomic analyses were performed using an Orbitrap Eclipse Tribrid mass spectrometer (Thermo Fisher Scientific) interfaced with an Easy-nLC 1200 ultra-high-performance liquid chromatography system (Thermo Fisher Scientific). Approximately 3 μg of peptides, resuspended in 5% acetonitrile (ACN) and 5% formic acid (FA), were injected onto either an in-house packed 100-μm inner diameter capillary column containing 35 cm of Accucore 150 resin (2.6 μm particle size, 150 Å pore size) or a PepMap EASY-Spray C18 analytical column (75 μm × 250 mm, 2 μm particle size, 100 Å pore size; Thermo Fisher Scientific). Peptides were separated over a 180-min gradient from 2% to 23% ACN in 0.125% FA at a flow rate of 500 nL/min. The electrospray voltage was maintained at 2.5 kV, and the ion transfer tube temperature was set to 300°C. Precursor ion separation was performed using either a FAIMS Pro or FAIMS Pro Duo interface (Thermo Fisher Scientific), operated under standard conditions with compensation voltages of −40 V, −60 V, and −80 V. For each compensation voltage, spectra were acquired in positive ion, data-dependent acquisition mode across an *m/z* range of 400–1400 using a 2-second duty cycle. Full MS scans were collected at a resolution of 120,000. Singly charged precursor ions were excluded from fragmentation, whereas multiply charged ions were selected for tandem mass spectrometry using standard automatic gain control (AGC) settings and higher-energy collisional dissociation (HCD) with a normalized collision energy (NCE) of 35%. A dynamic exclusion period of 30 seconds was applied. Quantification of tandem mass tag (TMT) reporter ions was performed using the multinotch synchronous precursor selection (SPS)-MS3 workflow, in which up to 10 SPS precursor ions were co-isolated and fragmented by HCD at 45% NCE. MS3 spectra were acquired in the Orbitrap at a resolution of 50,000 over an *m/z* range of 100–500. The mass spectrometry proteomics data have been deposited to the ProteomeXchange Consortium via the PRIDE partner repository with the dataset identifier PXD083672.

### Database searching

For CPT labeling optimization experiments, tandem mass spectra were searched using the Comet search engine against a UniProt protein sequence database containing both mouse (*Mus musculus*) and human (*Homo sapiens*) entries (downloaded in 2022). Reverse protein sequences were appended to generate a decoy database for false discovery rate (FDR) estimation, and common contaminant proteins, including human keratins and trypsin, were included in the search database. Database searches were performed using a precursor mass tolerance of 25 ppm and a fragment ion mass tolerance of 1.0 Da. Searches assumed fully tryptic digestion with a maximum of three missed cleavages. Variable modifications included methionine oxidation (+15.9949 Da) and cysteine CPT labeling (+221.08169 Da) for CPT-labeled cysteine proteome analyses. Peptide-spectrum matches were filtered using the target-decoy strategy to control the false discovery rate. Linear discriminant analysis (LDA) was applied to distinguish correct from incorrect peptide identifications, and peptide-level FDR was maintained below 0.5%. A minimum XCorr value of 1.0 was required, and peptide sequences consisting of six or fewer amino acids were excluded from downstream analyses.

### Calculating cysteine % oxidation

All TMT channels were normalized using a second ratio-check as described in Xiao et al., ^15^ to ensure equal peptide was loaded into each channel prior to enrichment, and that pipetting errors were corrected. For each site detected, the TMT reporter ion signal-to-noise ratio (S/N) from the oxidized channel was divided by the S/N from the total cysteine channel of the same protein to obtain the % cysteine oxidation value, controlling for changes in protein abundance. Eight biological replicates were used to calculate the average and standard error of the mean (SEM). Unless otherwise stated, we considered sites with percent oxidation changes ± 10% and P-value <0.05 as “changed significantly”.

### Analysis of CPT-proteomics data

All analyses were performed in R version 4.5.1 and GraphPad Prism version 11.0 unless otherwise noted. Protein subcellular locations were mapped to OxiMuscle from the Human Proteome Atlas based on validated data ^104^. Nucleoplasm, nuclear speckles, nuclear bodies, nuclear membrane, nucleus, nucleoli (fibrillar center), and nucleoli were consolidated into the nucleus annotation. To minimize localization ambiguity, only proteins with a single primary subcellular location reported were used. Secreted proteins were not present in the database, and therefore singly located proteins in ‘‘vesicles’’ category, and non-localized proteins were further queried from the secreted protein database downloaded from UniProt to generate a secreted protein list.

Protein-protein interaction networks (HCT and HEK) were downloaded from BioPlex 3.0, an affinity-purification/immunoprecipitation-MS-validated interactome built from 15,650 pull-downs^28^. The highest % oxidation value for each site was used to represent the maximum extent to which a protein was modified and mapped onto BioPlex 3.0. Network maps were visualized using Cytoscape 3.10.4^105^. Proteins with at least one site >10% or <-10% modified were regarded as highly modified proteins. To investigate whether the extent of coordinated redox modification in a protein network is more exercise-ubiquitous or exercise-selective, each network was scored by the percent of its constituent proteins that contain at least one highly oxidized cysteine in the same intervention. Non-identified proteins in communities were conservatively assigned as not highly oxidized to minimize false-positive results. Coefficient of variation (CV) is calculated for each community across all exercise types as an assessment of ubiquitous or exercise selective. Communities with no highly modified proteins in any tissues were labeled as non-redox regulated.

Gene Ontology (GO) Consortium pathways were downloaded from geneontology.org and mapped to the OxiMuscle data set by UniProt accession number to ensure only human sites matched. Pathways supported by literature related to exercise metabolism, skeletal muscle adaptive responses to exercise, and general biological function were utilized to rank each cysteine site (**Figure 3A**). All sites were initially filtered on having % oxidation modification >10 or <-10 and *P* < 0.05 compared to respective pre-exercise. After filtering, remaining sites were scored based on their appearance in the following exercise-adaptation -related GO pathways: glycolytic process, oxidative phosphorylation, protein biosynthetic process, response to exercise, human response to exercise (also known as human response to stress), autophagy, glucose homeostasis, glucose uptake, sarcomere, muscle contraction, muscle hypertrophy, translation, muscle adaptation, regulation of calcium signaling, mitophagy, muscle organ development, muscle cell differentiation, skeletal muscle tissue development, skeletal muscle atrophy, mitochondrion organization, regulation of mitochondrion organization, aerobic respiration, fatty acid beta-oxidation, glucose metabolic process, angiogenesis, regulation of NF-kB signaling, response to hypoxia, extracellular matrix organization, and response to reactive oxygen species. Sites were also scored separately by having appearance in the following cell biological pathways with importance for broad functioning: ubiquitin binding, regulation of protein phosphorylation, intracellular signal transduction, immune system process, cellular response to stress, cell differentiation, protein kinase binding, enzyme binding, cell proliferation, and inflammatory response. To determine the impact a modified cysteine has on a set of pathways, a cysteine impact score was determined by taking the sum of the absolute value of the % shift multiplied by the negative log10 *P*-value each time a site appeared in a pathway within the set.

### Cell culture

Mouse skeletal muscle cell line C2C12 myoblasts obtained from ATCC (Cat# CRL-1772), were cultured in DMEM (Corning #10-017-CV) without pyruvate, supplemented with 10% fetal bovine serum (GeminiBio #100-106) and 1% penicillin/streptomycin (P/S) (Corning #30-002-CI) at 37°C under a 5% CO_2_ atmosphere. All cells were washed with PBS (Corning #21-040-CV), detached using 0.25% trypsin (Gibco #25200-056), and subcultured every other day. When the cells reached 80-90% confluence, differentiation was induced by switching the growth medium to DMEM supplemented with 2% horse serum (Gibco #16050122) and 1% P/S (differentiation medium) for 7 days. The differentiation medium was changed every 24 h.

### CRISPR editing of p62C113S in C2C12 cell line

CRISPR/AsCas12a was used by the Initiative for Genome Editing and Neurodegeneration in the Department of Cell Biology at HMS to generate lines containing a T>A point mutation resulting in C113S amino acid substitution in the *Sqstm1* gene. The CRISPR target sequence was GGGGTGCCTCCTGAGCACATGGTG. The Ultramer sequence used to introduce the mutation was: 5’- CACCACAGGCCCGTTGCAACCATCACAGATCACATTGGGGTGCACCATGTT<u>G</u>CGGGGTGCCTCC TGAGCAC**t**TGGTGGGCGATGTTCCCGCCGGCACTCCTTCTTCTCTGCAGGGAGAGTCAAAGAGTC -3’ (point mutation lowercase; mutation in PAM region underscored). To create C2C12 cells harboring a homozygous p62/*Sqstm1* C113S mutation, 80 pmol of Alt-R CRISPR-Cas12a crRNA (IDT) was incubated with 63 pmol of AsCas12a protein for 10 minutes at room temperature and electroporated into 2 × 10^5^ C2C12 cells along with the Ultramer repair template and 39 pmol of Alt-R Cpf1 Electroporation Enhancer (IDT) using the Neon transfection system (Thermo Fisher Scientific). Mutants were identified by Illumina MiSeq sequencing. In addition, C113S mutation status was further confirmed by PCR amplification of genomic DNA, followed by Sanger sequencing using a primer set specific for *Sqstm1*. The primer sequences were as follows: forward, 5’- GGTAAAGGCTGAATCCGGGG-3’; reverse, 5’- GAGAGTGAGGGGTCAGGGTT-3’.

### EPS

Fully differentiated C2C12 myotubes in 4-well rectangular plates (Thermo Scientific #167063) or 12-well plates with #1.5 glass-like polymer coverslip bottom (Cellvis #P12-1.5P) were placed in a chamber for electrical stimulation (C-Dish; IonOptix, Westwood, MA) in an incubator at 37°C. Electrical stimulation was applied to the cells in the C-Dish using a C-Pace pulse generator (C-Pace EM; IonOptix). DMEM containing 2% horse serum was used during the EPS treatments. Cells in 4-well plates were treated with EPS at 1 Hz, 10 ms, 40 V for 2 hours. Cells in 12-well plates were stimulated at 1 Hz, 10 ms, 11 V for 2 hours.

### Transfection

C2C12 cells were seeded in 12-well plates, cultured for 24 hours and transfected pMRX-No-HaloTag7-mGFP-LC3 (Addgene #184901) with Lipofectamine™ 3000 Transfection Reagent (Invitrogen #L3000015) according to the manufacturer’s instructions.

### Live cell imaging

For intracellular H_2_O_2_ imaging, we followed previously described protocols ^69,106^. In brief, cultured myotubes transduced with adenoviruses expressing HyPer3-DAAO or HyPer7-DAAO were excited at 420 and 490 nm, and emission was captured at 510-530 nm using an Olympus IX81 microscope equipped with a 20 × oil immersion objective (PlanSApo, Olympus) and an ImagEM CCD camera (Hamamatsu, Bridgewater, NJ, USA) at a binning of 4. Data were acquired using MetaFluor Software (Molecular Devices, San Jose, CA, USA) and intracellular H_2_O_2_ was quantified after background subtraction by calculating the ratio of R, where R represents the ratio of the 490 nm to the 420 nm signal.

### Immunofluorescence

For the detection of endogenous p62 and LC3 puncta, C2C12 myoblast cells were seeded in 12-well plates (Cellvis #P12-1.5P). When cells were around 60% confluent, plasmid HyPer7.2DAAO-NES were transfected with Lipofectamine 3000 as described above. Forty-eight hours after transfection, growth medium was switched to medium containing 4 mM D-alanine for 2 hours. After treatment, cells were fixed in 4% formaldehyde in PBS for 10 min at room temperature. Cells were permeabilised with 0.5% Triton X-100 for 5 min at room temperature. Cells were blocked for one hour in 5% normal goat or rabbit serum in PBS with 0.05% Tween-20. Cells were incubated with primary antibodies overnight at 4 °C. Primary antibodies used in this study include guinea pig anti-p62 (Progen #Gp62-C, 1:500) and rabbit anti-LC3 (CST #3868, 1:500). Cells were washed three times and incubated with goat anti-guinea pig IgG secondary antibody conjugated with Alexa Fluor 647 (Life Technologies, Cat# A21450, 1:5000) or anti-rabbit IgG conjugated with Alexa Fluor™ 568 (Invitrogen #A-11004, 1:5000) for 1 h at room temperature.

For basal autophagic flux assays in C2C12 myoblasts, cells were seeded onto a 12-well glass-bottom dish (Cellvis #P12-1.5P). After 24 hours, plasmid pMRX-No-HaloTag7-mGFP-LC3 (Addgene #184901) was transfected with Lipofectamin 3000 as described above. On the third day, cells were incubated in growth medium with 100 nM TMR-conjugated Halo ligand (Promega, G8251) for 20 min and washed with PBS twice. The cells were then fixed in 4% formaldehyde in PBS for 10 min at room temperature.

For basal and EPS autophagic flux assays in C2C12 myotube, wild type and C113S cells were seeded onto a 12-well glass-bottom dish (Cellvis #P12-1.5P). On the second day, plasmid pMRX-No-HaloTag7-mGFP-LC3 (Addgene #184901) was transfected with Lipofectamin 3000. When cells reached to 80-90% confluence, growth medium was switched to 2% horse serum medium to induce myotubes. Seven days after differentiation, cells were incubated in differentiation medium with 100 nM TMR-conjugated Halo ligand (Promega, G8251) for 20 min and then EPS was applied to cells at 1 Hz, 10 ms, 11 V for 2 hours. Cells were then fixed in 4% formaldehyde in PBS for 10 min at room temperature. After treatment, cells were washed and mounted with ProLong Glass Antifade Mountant with NucBlue Stain (Invitrogen #P36981) and imaged with an LSM 980 with Airyscan 2 confocal microscope (Zeiss) using a 100× Plan-Apo/1.4 NA Oil objective. Images were processed with Zeiss ZEN Blue software and analyzed using ImageJ software (NIH). For double-positive puncta (p62^+^LC3^+^) identification, the positions of the puncta were identified with Find Maxima and marked as points on a binary image for both fluorescence channels. With the binarized images, double-positive puncta were identified with the AND function in Image Calculator (i.e., ‘p62’ image AND ‘LC3’ image). The resulting puncta after each operation were counted with Analyze Particles.

### CRISPR editing of p62C113S in C57BL/6J mice

The p62C113S mouse was generated by CRISPR/Cas9-mediated homology-directed repair at the murine Sqstm1 locus on a C57BL/6J background. A single guide RNA (sgRNA) targeting exon 2 proximal to codon 113 (5′-ACATCGCCCACCATGTGCTC-3′) was designed to introduce a cysteine-to-serine substitution (TGT→AGT). An Alt-R™ single-stranded HDR donor oligonucleotide (Integrated DNA Technologies) containing the desired C113S mutation and silent nucleotide substitutions to disrupt Cas9 re-cutting was synthesized with ∼70-bp homology arms flanking the editing site (5′- TAAGTTCTGGATGGACTCTTTGACTCTCCCTGCAGAGAAGAAGGAGTGCCGGCGGGAACATCGC CCACCA**a**GTGCTCA<u>a</u>GAGGCACCCCGAAACATGGTGCACCCCAATGTGATCTGTGATGGTTGCAA CGGGCCTGTG-3′ (point mutations lowercase; PAM region underscored)). Recombinant Cas9 protein (Integrated DNA Technologies, #1081058) was complexed with synthetic sgRNA to form ribonucleoprotein (RNP) complexes, which were co-injected with the HDR donor oligo into the pronuclei of fertilized one-cell embryos at the Dana-Farber/Harvard Cancer Center (DF/HCC) Transgenic Mouse Core. Injected zygotes were cultured to the two-cell stage and transferred into pseudopregnant recipient females. Founder pups were screened by PCR amplification of the targeted region followed by Sanger sequencing (Azenta Life Sciences/Genewiz, Burlington, MA) to confirm precise incorporation of the C113S substitution without additional indels. The primer sequences were as follows: forward, 5’-GGTAAAGGCTGAATCCGGGG-3’; reverse, 5’- GAGAGTGAGGGGTCAGGGTT-3’. Correctly targeted founders were backcrossed to C57BL/6J mice to establish germline transmission, and heterozygous offspring were intercrossed to obtain homozygous p62C113S mice.

### Study approval

All animal experiments were carried out under NIH guidelines for the care and use of laboratory animals, and all animal protocols were approved by the Dana-Farber Cancer Institute Institutional Animal Care and Use Committee (protocol 24-029).

### Indirect calorimetry measurement via Promethion Core system

The Promethion Core CGF system (Sable, North Las Vegas, NV) was used to perform indirect calorimetry (IC) measurements. The system is equipped with a baseline chamber to measure the oxygen consumption (VO_2_), carbon dioxide production (VCO_2_) and energy expenditure. Food and water intake, body weight and movement were monitored simultaneously.

### Grip strength test

To measure grip strength, a computerized grip strength meter (Bio-GS4, BIOSEB, Vitrolles, France) was used. Each mouse was held gently by the base of the tail and lowered toward the grid mesh of the apparatus until its paws firmly grasped the pull bar. The mouse was then pulled horizontally backward along the axis of the sensor with a steady, smooth and continuous motion until its grip was released. The digital force transducer recorded the peak resistance force in newtons. Each mouse underwent four consecutive trials with an inter-trial rest interval of approximately 15 seconds to prevent muscle fatigue. The average of the four independent measurements was calculated and used for statistical analysis.

### Treadmill acclimation and exercise capacity test

Mice were acclimated to treadmill running for 3 consecutive days prior to the exercise capacity test using a motorized treadmill (Columbus Instruments, OH, USA). During each acclimation session, mice ran at a constant speed of 5 m/min for 6 min. The treadmill was initially maintained at a 0° incline for 2 min, followed by a gradual increase to a 5° incline over 1 min. Mice then ran at a 5° incline for 2 min, after which the incline was gradually returned to 0° over 1 min. Exercise capacity was assessed using an incremental treadmill test. Mice initially ran at 5 m/min for 2 min, after which the speed was increased to 8 m/min and subsequently increased by 2 m/min every 2 min (10, 12, 14, 16, 18, 20, 22, and 24 m/min) until exhaustion. Exhaustion was defined as the inability of a mouse to continue running and remaining on the electric shock grid for at least 5 s without resuming running. The start and exhaustion times were recorded for each mouse and used to determine total running time, while total running distance was recorded directly by the treadmill software.

### SP3-based sample preparation for quantitative proteomics

Skeletal muscle samples were processed using a single-pot solid-phase-enhanced sample preparation (SP3) workflow prior to LC–MS/MS analysis. Soleus and gastrocnemius muscles were lysed in 50 mM HEPES buffer (pH 8.5) containing 2% SDS and protease inhibitors (Roche). Samples were homogenized using a bead beater in a cold room and further sonicated to shear nucleic acids and reduce viscosity. Lysates were centrifuged at 21,000 × g for 15 min at 4°C to remove insoluble debris, and protein concentration was determined using a bicinchoninic acid (BCA) assay. For proteomic sample preparation, 10 μg of total protein was adjusted to 45 μL in lysis buffer containing 1% SDS. Disulfide bonds were reduced with 5 mM dithiothreitol (DTT) at 37°C for 30 min, followed by alkylation with 14 mM iodoacetamide at room temperature for 45 min in the dark. SP3 magnetic beads were prepared by combining hydrophilic and hydrophobic Sera-Mag SpeedBeads and washing twice with water before use. One microliter of SP3 bead slurry was added to each sample, followed by ethanol to a final concentration of 50% to induce protein binding to the beads. Samples were incubated at room temperature for 5 min and beads were immobilized using a magnetic rack. Beads were washed three times with 80% ethanol to remove contaminants and residual detergent. Proteins bound to SP3 beads were digested overnight at 37°C in 50 mM ammonium bicarbonate containing sequencing-grade trypsin (0.5 μg) and LysC (0.3 μg). Following digestion, peptide-containing supernatants were separated from beads using a magnetic rack and acidified with formic acid prior to desalting. Peptides were desalted using homemade C18 StageTips packed with C18 membrane disks. StageTips were conditioned sequentially with acetonitrile, 70% acetonitrile/1% formic acid, and 1% formic acid before sample loading. Bound peptides were washed with 1% formic acid and eluted with 40% and 70% acetonitrile containing 1% formic acid. Eluted peptides were dried in a SpeedVac concentrator and resuspended in 0.1% formic acid. Samples were centrifuged at maximum speed for 10 min, and the clarified peptide supernatant was loaded onto Evotips for LC–MS/MS analysis on the timsTOF (Bruker) mass spectrometer.

Peptides were separated using a PepSep C18 Column (15 cm × 150 μm, particle size 1.5 μm) and a 30 SPD method (44-min gradient) on Evosep One. CaptiveSpray source was operated at 1600 V with a dry gas flow of 3 L/min at 180 °C. The mass spectrometer was operated in data-independent acquisition with parallel accumulation serial fragmentation (DIA-PASEF) mode. MS parameters: polarity positive, scan m/z range 100–1700, mobility (1/K0) range 0.60–1.60 V⋅s/cm^2^, ramp time 75 ms, accumulation time 75 ms. MS/MS was performed using optimized window settings with 1.7 s cycle time estimate, 1 MS1 ramp, 20 MS/MS ramps, 60 MS/MS windows, 350.7–1250.6 Da mass range, and 0.6–1.45 V⋅s/cm2 mobility range (1/K0). Linear mobility-dependent collision energy was applied with 20 eV at 0.60 V⋅s/cm2 and 59 eV at 1.6 V⋅s/cm^2^. Data analysis was performed using DIA-NN (version 2.3.0) with developer-recommended settings (default except using 15 ppm mass errors). In silico predicted library was generated from a FASTA file containing 54,707 Mus musculus (mouse) protein sequences retrieved from UniProt (https://www.uniprot.org/proteomes/UP000000589). The mass spectrometry proteomics data have been deposited to the ProteomeXchange Consortium via the PRIDE partner repository with the dataset identifier PXD083723.

### Immunofluorescence for muscle fiber characterization

Frozen OCT-embedded plantaris muscles were sectioned at 10 μm on a Leica cryostat. Immunofluorescently stained slides were observed with a fully automated Nikon Ti2 widefield light microscope with the 20 × objective lens. Fluorescent images were taken using a high-sensitivity Hamamatsu Flash 4.0 LT camera. To assess myofiber cross-sectional area (CSA) and fiber type^84^, muscle sections were blocked for 1 h with mouse immunoglobulin G (IgG) blocker, MOM (Vector, San Diego, CA, USA cat# MKB-2213). Sections were then incubated in primary laminin antibody (1:200, Sigma, St. Louis, MO, USA, cat# L9393) and primary antibodies BA-D5, SC-71, and BF-F3 (1:100 Developmental Studies Hybridoma Bank, Univ. of Iowa) overnight at 4°C, followed by secondary antibody Alexa Fluor 350 (Invitrogen, cat# A11046) for laminin and Alexa Fluor 647, 555, and 488 for fiber types (Invitrogen, cat# A21242, A21121, and A21426). CSA and fiber type were measured using semi-automatic muscle analysis with segmentation of histology, a MATLAB application (SMASH) and confirmed with ImageJ software. Using the SMASH software, an average of 536 ± 60 SEM plantaris fibers were included in the CSA analysis for each muscle.

### Western blot

For western blot analysis, mouse gastrocnemius and soleus were lysed in 2% SDS in PBS buffer supplemented with EDTA-free cOmplete protease inhibitor (Roche) then sonicated to shear chromatin using tip sonicator at 20% amplitude for 15 seconds (Qsonica). The protein concentration was determined using a Pierce™ BCA assay kit (Thermo Fisher Scientific, USA). Lysate was adjusted to equal protein concentration across samples and diluted with 4x NuPAGE LDS sample buffer (Thermo Fisher Scientific, USA) and samples were heated for 10 min at 95°C. Samples were separated on 4 –12% NuPAGE BisTris (Thermo Scientific, USA) gels using MOPS running buffer (Thermo Fisher Scientific, USA). Proteins were transferred to PVDF membranes using the iBLOT2 transfer system (Thermo Fisher Scientific, USA) with iBLOT2 PVDF transfer stacks (Thermo Fisher Scientific, USA). Membranes were blocked with 3% BSA (Sigma, USA) in TBS + 0.1% Tween-20 (Boston BioProducts, USA). Membranes were incubated with primary antibodies overnight at 4°C. Membranes were washed three times with TBS + 0.1% Tween-20 (Boston BioProducts, USA) followed by incubation with secondary antibodies for 1 hour at room temperature in TBS + 0.1% Tween-20 (Boston BioProducts, USA) containing 3% BSA (Sigma, USA). Membranes were then washed three times with TBS + 0.1% Tween-20 (Boston BioProducts, USA).

### Statistical analysis

All experiments were performed at least three times unless otherwise noted. Mean values for individual experiments are expressed as means ± standard error mean (SEM). Distributions were analyzed for normality with the Shapiro– Wilk test. Comparisons between two groups were assessed by two-sided Student’s t-test (paired or unpaired, as appropriate) if normally distributed, by Welch’s t-test where unequal variances were assumed or by Mann–Whitney U test if not normally distributed. Comparisons involving two independent factors (e.g., genotype × treatment or genotype × exercise) were assessed by two-way ANOVA with Tukey’s post-hoc test. Repeated measures across a single factor over time were assessed by one-way repeated-measures ANOVA with Tukey’s post hoc test. Network enrichment analyses were assessed by Fisher’s exact test, with P-values corrected for multiple comparisons using the Benjamini-Hochberg false discovery rate (FDR) procedure; a significance threshold of FDR-adjusted P < 0.05 was applied. A *P*-value of < 0.05 was considered statistically significant. All physiological and imaging studies were performed and analyzed by scientists blinded to genotype and treatment. Statistical analyses were performed using GraphPad Prism 11.0 (GraphPad Software, La Jolla, CA) and R (version 4.5.1; RStudio, Boston, MA).

#### AI-assisted code development

R code used for data analysis and visualization was developed and debugged with assistance from ChatGPT (OpenAI) and Claude (Anthropic). The analytical approaches were determined by the authors, and all AI-assisted code and resulting analyses were reviewed and validated by the authors.

## ACKNOWLEDGEMENTS

The project was funded by a Synergy Grant from the Novo Nordisk Foundation (Grant no. NNF20OC0063709). This work was also supported by the Claudia Adams Barr Program, the Lavine Family Fund, the Pew Charitable Trusts, NIH DK123095, NIH AG071966, the Mark Foundation, the Smith Family Foundation, and the American Federation for Aging Research, all awarded to E.T.C. E.T.C. is an HHMI investigator. Y.L. was supported by Postdoctoral Fellowships from the Swedish Society of Medicine (Grant no. PD20-0102) and the American Heart Association (Grant no. 25POST1376679). C.H.-O. was supported by a postdoctoral grant from the Danish Diabetes Academy, funded by the Novo Nordisk Foundation (Grant no. NNF17SA0031406). We thank HMS Transgenic Mouse Core and The Initiative for Genome Editing and Neurodegeneration in the Department of Cell Biology at HMS for their support.

## AUTHOR CONTRIBUTIONS

Y.L., J.J.P., E.A.R. and E.T.C. carried out study conceptualization, design and direct research. Y.L. and J.J.P. designed and performed experiments, analyzed data, interpreted results, and prepared figures. A.R. and N.B. assisted with Redox proteomics experiment and data analysis. C.H.-O. contributed to the design, imaging and quantification of myoblast autophagic flux experiments. S.S. and B.Z. assisted with general and Redox proteomics. T.C. and M.W.-W. assisted with ROS measurement. J.Z. assisted with gene editing. C.H.-O., T.J., C.T.V., C.S.C. and B.K. contributed to human exercise coordination, sample collection and data discussion. S.M.W., J.J.P. and Y.L. built the website. The manuscript was written by E.T.C., Y.L. and J.J.P. with the help of all authors.

## CONFLICTS OF INTEREST

E.T.C. is a co-founder, equity holder, and board member of Matchpoint Therapeutics, a co-founder and equity holder in Aevum Therapeutics, and paid consultant for WndrHLTH.

### Declaration of generative AI and AI-assisted technologies in the manuscript preparation process

During the preparation of this work, the authors used Claude (Anthropic) to proof read the language and readability of the manuscript. After using this tool, the authors reviewed and edited the content as needed and take full responsibility for the content of the publication.

## SUPPLEMENTARY MATERIAL

**Figure S1.**
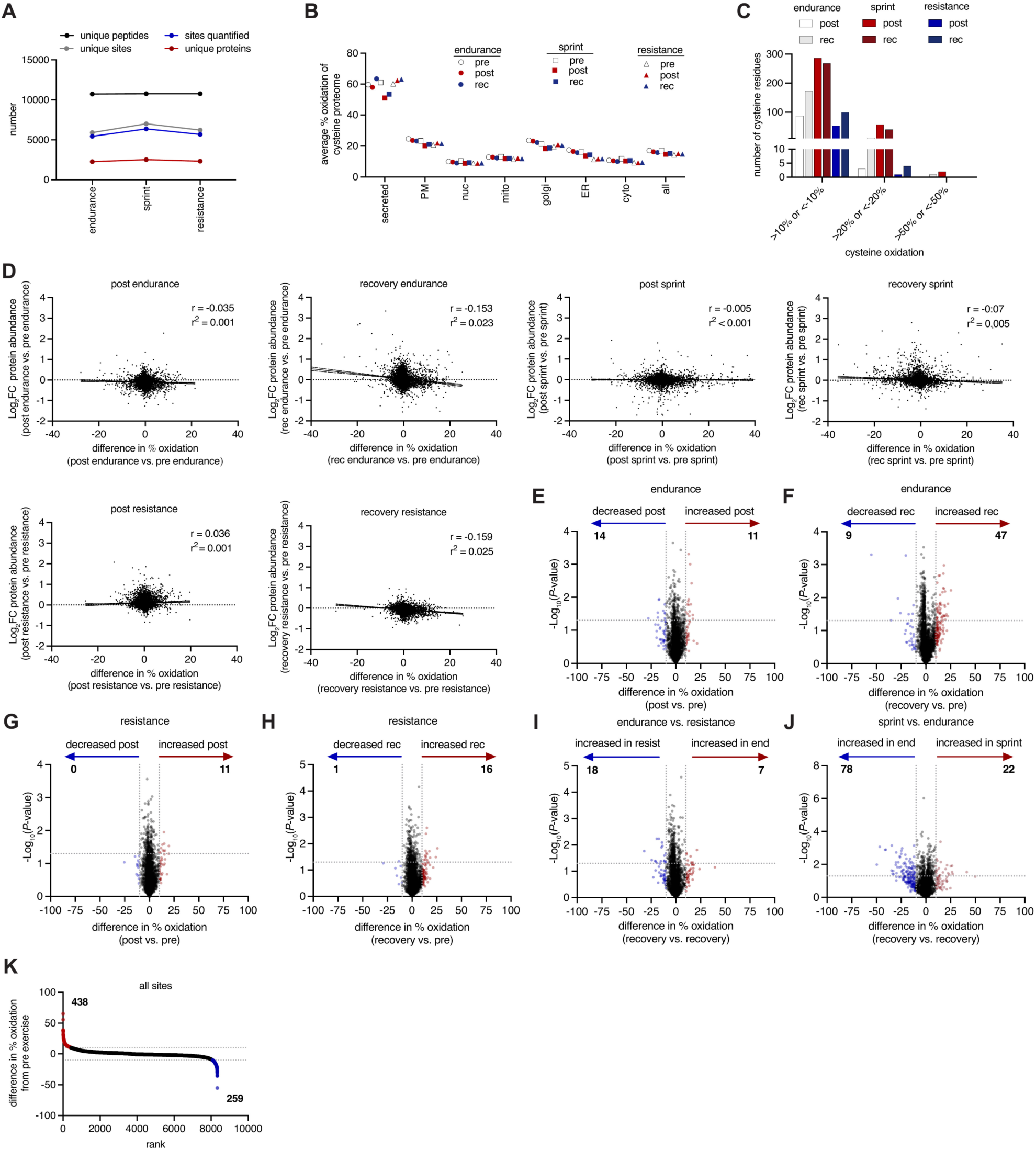
OxiMuscle Data Characteristics. **(A)** Numbers of unique peptides, cysteine sites, cysteine sites quantified and unique proteins identified. [Values are sums across n = 8 biological replicates per exercise condition.] **(B)** Average percent oxidation distribution of the cysteine proteome within subcellular locations for each exercise type. **(C)** Proportion of cysteines with a shift in oxidation state from respective pre-exercise of > 10% or < -10%, > 20% or < - 20%, and > 50% or < -50%, in each exercise type, post and in recovery. **(D)** Correlation between log₂ fold change in total protein abundance and percent shift in cysteine oxidation state, both calculated relative to respective pre-exercise levels at post-exercise and recovery timepoints, across all exercise types. Total cysteine channel was used to represent changes in total protein abundance measured. **(E-F)** Pairwise comparison of cysteine modification state between endurance post and recovery vs. the pre-exercise proteome. [n = 8 biological replicates per condition. Differentially modified cysteines were identified by two-sided paired Student’s t-test; highlighted/colored cysteines indicate sites reaching |Δ% modification| > 10%.] **(G-H)** Pairwise comparison of cysteine modification state between resistance post and recovery vs. the pre-exercise proteome. [n = 8 biological replicates per condition; statistical test and thresholds as in (E-F).] **(I)** Pairwise comparison of cysteine modification state between endurance recovery vs. resistance recovery proteome. [n = 8 biological replicates per condition; statistical test and thresholds as in (E-F).] **(J)** Pairwise comparison of cysteine modification state between sprint recovery vs. endurance recovery proteome. [n = 8 biological replicates per condition; statistical test and thresholds as in (E-F).] **(K)** Ranking all cysteine sites quantified by percent shift from respective pre-exercise. All analyses use n = 8 biological replicates (participants) per exercise condition. Statistical significance of cysteine oxidation changes was assessed by two-sided paired Student’s t-test; cysteines passing P < 0.05 are annotated as significantly modified, cysteines passing P < 0.05 and |Δ% modification| > 10% are annotated as highly modified.

**Figure S2.**
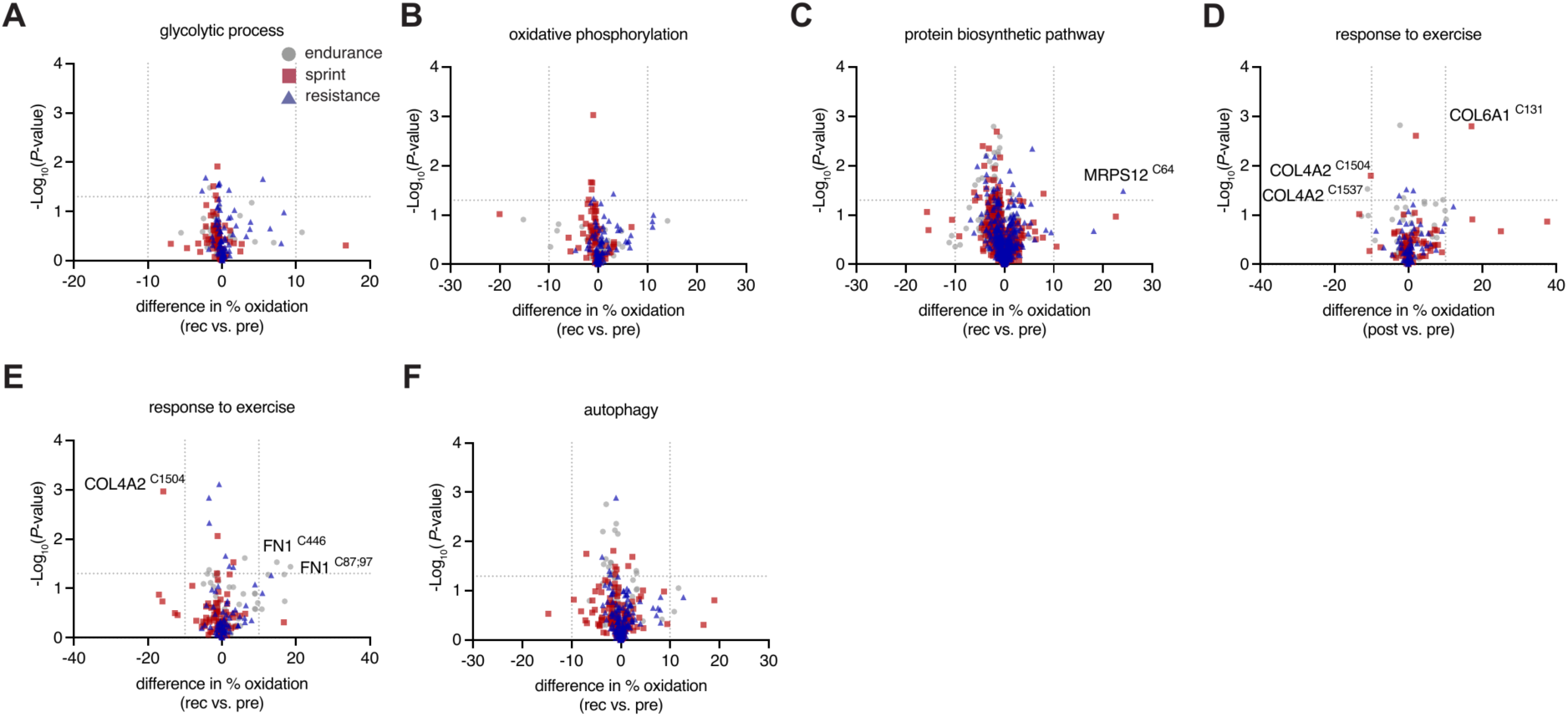
Exercise-modality-specific Cysteine Oxidation Changes in Proteins Across Key Metabolic and Structural Pathways. **(A)** Pairwise comparison of cysteine modification state recovery vs. pre-exercise proteome in each exercise type and filtered for proteins in the glycolytic process GO pathway. [n = 8 biological replicates per exercise condition; highlighted cysteines reach P < 0.05 (two-sided paired Student’s t-test) and |Δ% modification| > 10%.] **(B)** Pairwise comparison of cysteine modification state recovery vs. pre-exercise proteome in each exercise type and filtered for proteins in the oxidative phosphorylation GO pathway. [n = 8 biological replicates per condition; statistical test and thresholds as in (A).] **(C)** Pairwise comparison of cysteine modification state recovery vs. pre-exercise proteome in each exercise type and filtered for proteins in the protein biosynthetic GO pathway. MRPS12, small ribosomal subunit protein uS12m. [n = 8 biological replicates per condition; statistical test and thresholds as in (A).] **(D)** Pairwise comparison of cysteine modification state post vs. pre-exercise proteome in each exercise type and filtered for proteins in the response to exercise GO pathway. FN1, fibronectin; COL4A2, collagen alpha-2(IV) chain. [n = 8 biological replicates per condition; statistical test and thresholds as in (A).] **(E)** Pairwise comparison of cysteine modification state recovery vs. pre-exercise proteome in each exercise type and filtered for proteins in the response to exercise GO pathway. COL6A1, collagen alpha-1(VI) chain. [n = 8 biological replicates per condition; statistical test and thresholds as in (A).] **(F)** Pairwise comparison of cysteine modification state recovery vs. pre-exercise proteome in each exercise type and filtered for proteins in the autophagy GO pathway. [n = 8 biological replicates per condition; statistical test and thresholds as in (A).] All analyses use n = 8 biological replicates (participants) per exercise condition. Highlighted cysteines pass P < 0.05 (two-sided paired Student’s t-tests) and |Δ% modification| > 10%.

**Figure S3.**
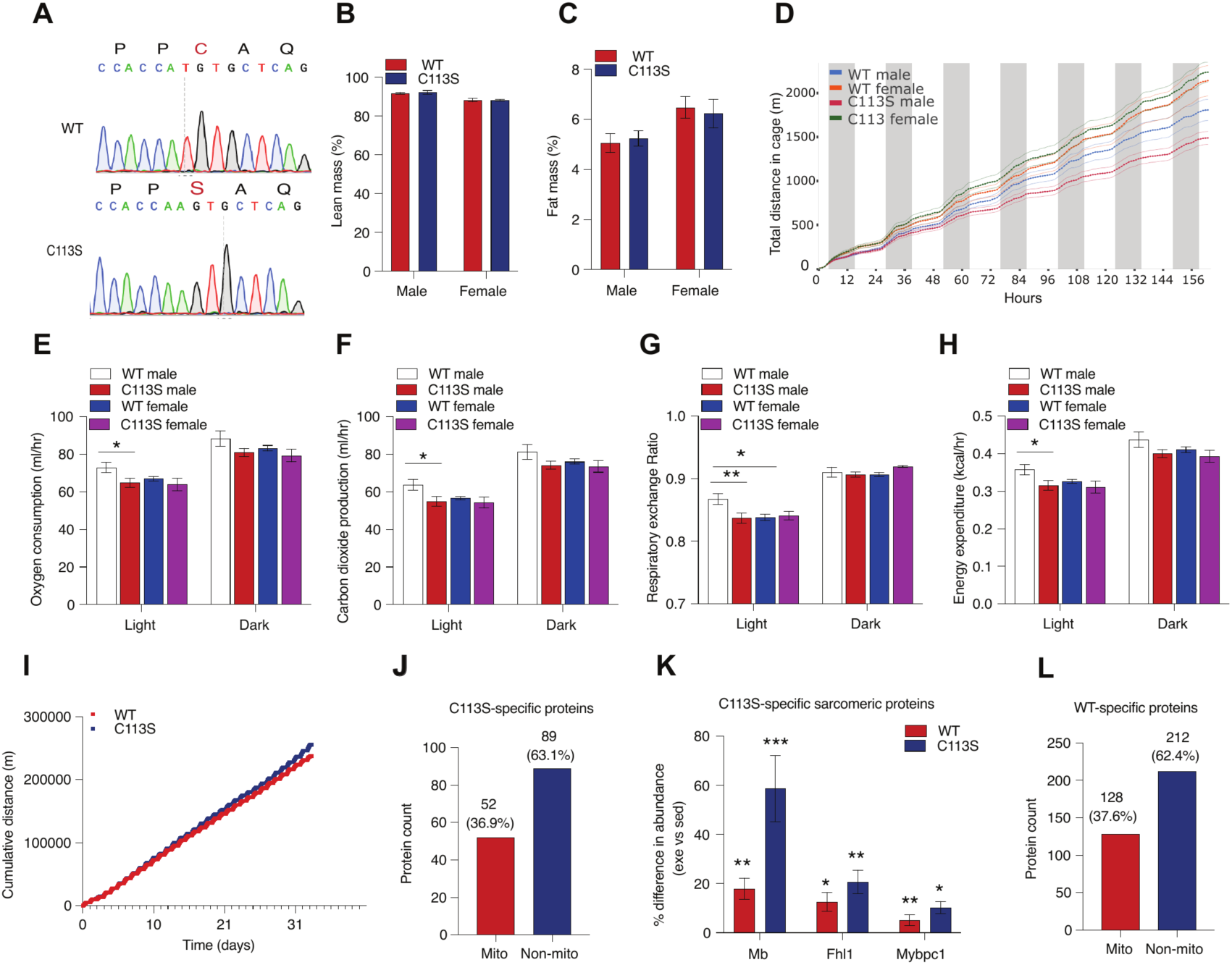
Metabolic Characterization of p62 C113S Mice at Baseline and Proteome Remodeling After Exercise. **(A)** Sanger sequencing chromatograms of C2C12 cells confirming the CRISPR/Cas9-mediated TGT→AGT (C113S) substitution. [Chromatogram representative of n = 3 independently isolated clonal lines.] **(B–C)** Body composition analysis showing lean mass and fat mass percentages in male and female WT and p62 C113S mice measured by EchoMRI. [n = 9–11 per group.] **(D)** Total distance in cage (m) assessed under basal conditions. [n = 4–7 per group.] **(E–H)** Indirect calorimetry measurements including oxygen consumption (VO₂), carbon dioxide production (VCO₂), respiratory exchange ratio (RER), and energy expenditure in WT and p62 C113S mice. [n = 4–7 per group.] **(I)** Cumulative running distance (m) of WT and C113S mice during voluntary wheel running. [n = 8 per group.] **(J)** Proportion of mitochondrial and non-mitochondrial proteins among C113S-specific exercise-responsive proteins. [n = 9–11 per group.] **(K)** Differential regulation of sarcomeric proteins between WT and p62 C113S mice following exercise. [n = 9–11 per group.] **(L)** Proportion of mitochondrial and non-mitochondrial proteins among WT-specific exercise-responsive proteins. [n = 9– 11 per group.] For **(B-C, E-H)**, data are presented as mean ± SEM. Statistical significance was determined using two-way ANOVA with Tukey’s post-hoc test. For **(K)**, bars show mean percent difference in abundance (exercise vs. sedentary) ± SEM per genotype. Statistical significance for each protein was determined by a two-sample Welch’s t-test comparing individual exercised versus sedentary mice within each genotype; *P < 0.05, **P < 0.01, ***P < 0.001.

