## Supplementary material for "A quantitative redox proteome of the human muscle response to exercise": Table S1

**Table S1. Subject characteristics**

| **Characteristic** | **Mean ± SEM** |
| --- | --- |
| Men, n | 8 |
| Age, years | 26.3 ± 1.6 |
| Weight, kg | 76.1 ± 3.9 |
| BMI, kg/m² | 23.5 ± 0.7 |
| **DXA** |  |
| Fat, % | 21.2 ± 2.0 |
| Lean mass, kg | 57.0 ± 1.8 |
| **Fitness level** |  |
| VO₂peak, mL·min⁻¹ | 3236 ± 172 |
| VO₂peak, mL·min⁻¹·kg⁻¹ | 42.6 ± 1.5 |
| Maks watt completed, W | 278 ± 16.0 |
| **Oxygen consumption and energy expenditureᵃ** |  |
| **Endurance (n = 8)** |  |
| VO₂, mL/min | 1975 ± 103 |
| Energy expenditure, kJ | 3609 ± 189 |
| **Sprint (n = 5)ᵇ** |  |
| VO₂, mL/min | 2001 ± 106 |
| Energy expenditure, kJ | 382 ± 16 |
| **Resistance (n = 3)ᵇ** |  |
| VO₂, mL/min | 779 ± 1.3 |
| Energy expenditure, kJ | 198 ± 0.3 |

*Data are mean ± SEM.*

*^a^ Energy expenditure was calculated from oxygen uptake in a subset of trials. For endurance and resistance, oxygen consumption was measured by indirect calorimetry; for sprint, oxygen consumption was estimated from the relationship between heart rate and oxygen consumption established during the peak VO₂ test. Values reflect the entire exercise period, including rest intervals between bouts for sprint and resistance.*

*^b^ Sprint and resistance oxygen-consumption/energy-expenditure measurements were available for a subset of the n = 8 participants; all other measures reflect the full cohort (n = 8).*
