## Supplementary material for "A quantitative redox proteome of the human muscle response to exercise": Table S5

**Supplementary Table S5. Genotype distribution of offspring from p62 C113S heterozygous crosses**

|  | **WT** | **Heterozygous p62 C113S** | **Homozygous p62 C113S** | **Total** |
| --- | --- | --- | --- | --- |
| **Observed N** | 305 (173F + 132M) | 522 (259F + 263M) | 160 (85F + 75M) | **987** |
| **Observed %** | 30.9% | 52.9% | 16.2% | **100%** |
| **Expected N (1:2:1 ratio)** | 246.75 | 493.5 | 246.75 | **987** |
| **Expected %** | 25% | 50% | 25% | **100%** |

**Chi-square goodness-of-fit test (observed vs. expected 1:2:1 Mendelian ratio):** χ² = 45.90, df = 2, p = 1.1 × 10⁻¹⁰. The observed genotype distribution deviated significantly from the expected 1:2:1 Mendelian ratio, reflecting a reduced proportion of homozygous p62 C113S offspring (16.2% observed vs. 25% expected) and a corresponding excess of WT offspring (30.9% observed vs. 25% expected), consistent with partial embryonic or perinatal lethality of the homozygous genotype.
